# PaRXLR40, a broad cell death suppressor of the kauri dieback pathogen *Phytophthora agathidicida*, targets a plant ARM/BTB domain-containing protein

**DOI:** 10.1101/2025.11.05.686655

**Authors:** Mariana Tarallo, Yanan Guo, Hazel McLellan, Ellie L. Bradley, Rosie E. Bradshaw, Petra C. Boevink, Paul R. J. Birch, Carl H. Mesarich

## Abstract

- *Phytophthora agathidicida*, the causal agent of kauri dieback, secretes RXLR effector proteins to promote host colonisation. One of these, PaRXLR40, was previously shown to suppress immune responses in *Nicotiana benthamiana*, but its mechanism of action and contribution to virulence remained unclear.
- To investigate PaRXLR40 function, we used comparative approaches in *N. benthamiana* and *Agathis australis* (kauri), including RNA interference (RNAi), transient expression assays, confocal microscopy, yeast two-hybrid screens, and infection assays. We also examined host protein interactors and tested mutant variants to evaluate functional domains.
- Silencing *PaRXLR40* reduced *P. agathidicida* colonization in *N. benthamiana* and *A. australis*. PaRXLR40 interacted with a host BTB/ARM domain protein (ARIA), previously implicated in abscisic acid (ABA) signalling. ARIA suppressed immunity and promoted infection, while interacting with NbSOG1, a DNA damage-associated transcription factor that enhanced resistance when overexpressed. External application of ABA enhanced *P. agathidicida* infection in both hosts, supporting the hypothesis that PaRXLR40 may hijack host ABA signalling through ARIA to promote susceptibility.
- Our findings show that PaRXLR40 targets ARIA to manipulate host immunity and promote virulence. The interaction between ARIA and SOG1 suggests PaRXLR40 may interfere with host transcriptional reprogramming. PaRXLR40 represents a potential target for future RNAi-based strategies to reduce kauri dieback.

## 1 Introduction

Pathogenic oomycetes cause some of the most devastating diseases of plants (Kamoun *et al*., 2015; Fones *et al*., 2020). Examples include the soybean stem and root rot pathogen, *Phytophthora sojae*, as well as the sudden oak death pathogen, *Phytophthora ramorum* (Günwald *et al*., 2008). The diseases caused by these types of oomycetes cost the agriculture and forestry industries billions of dollars per year worldwide, but others also have severe environmental and social impacts. One of these is root and collar dieback of New Zealand (NZ) kauri (*Agathis australis*), hereafter referred to as kauri dieback, which is caused by the clade 5 soil-borne oomycete, *Phytophthora agathidicida* (Beever *et al*., 2009). Protecting *A. australis* from this disease is of utmost importance, since this tree species is one of the world’s largest ancient Araucariaceae conifers and has immense cultural significance for NZ Māori (Lambert *et al*., 2018). Currently, the management of kauri dieback is predominantly reliant on containment to avoid pathogen spread and the injection of phosphite into infected trees to minimise effects of the disease (Horner *et al*., 2015; Bradshaw *et al*., 2020). However, preliminary screening assays suggest there is natural tolerance to kauri dieback in the *A. australis* population that could potentially be harnessed to assist with the long-term management of this disease (Bradshaw *et al*., 2020).

Understanding how pathogens and their hosts interact at the molecular level provides important knowledge for breeding and selecting plants with resistance or tolerance against disease (Sugimoto *et al*., 2012; Vleeshouwers & Oliver, 2014). This resistance or tolerance is often based on their multilayered immune system made up of both extracellular (cell surface) and intracellular immune receptors that, upon recognition of pathogen- or damage-associated molecules, crosstalk with each other and hormone signalling pathways to deliver robust defence responses (Ngou *et al*., 2022). Successful pathogens, however, secrete effector proteins to suppress host immunity and promote infection (Lo Presti *et al*., 2015; Rocafort *et al*., 2020). This is achieved through the manipulation of diverse host targets, with a subset shown to manipulate, for example, transcription factors and hormone signalling components (Anderson *et al*., 2015; He *et al*., 2020). Among plant hormones, salicylic acid (SA) and jasmonic acid (JA) are typically associated with immune activation, whereas abscisic acid (ABA) has a more complex role. In several plant-pathogen interactions, ABA can promote susceptibility by interfering with immune signalling (Singh & Roychoudhury, 2023).

Like other *Phytophthora* pathogens, *P. agathidicida* secretes RXLR effectors during infection of *A. australis*. These effectors typically carry an RXLR motif (Arginine-any amino acid-Leucine-Arginine) located approximately 25-60 amino acids downstream of the signal peptide and are often immediately followed by an EER motif (Glutamate-Glutamate-Arginine) (Whisson *et al*., 2007; Wang *et al*., 2023). In addition to these motifs, many RXLR effectors possess a WY domain, a structural fold stabilized by conserved tryptophan (W) and tyrosine (Y) residues in the C-terminal region. This fold has been shown to contribute to effector stability and host target interactions during infection (Boutemy *et al*., 2011). Many *Phytophthora* RXLR effectors have a role in virulence by targeting different host molecules to suppress host immunity (Bos *et al*., 2010; Yang *et al*., 2019; Mach, 2021; Wang *et al*., 2023), while the recognition of other *Phytophthora* RXLR effectors by plant immune receptors triggers defence responses (He *et al*., 2020). A recent study identified 147 RXLR genes in the *P. agathidicida* genome (Cox *et al*., 2022). We previously showed that nine of the effectors encoded by these genes can interact with the immune system of angiosperms, specifically of *Nicotiana* spp. (Guo *et al*., 2020). Although *P. agathidicida* is a pathogen of gymnosperms, *Nicotiana benthamiana* is an alternative host in the laboratory, making it a valuable model for functional assays involving effectors from this pathogen (Bradley, 2022). Among the effectors of *P. agathidicida* that have been studied in *N. benthamiana* are PaRXLR24 and PaRXLR40, which are phylogenetically closely related to each other and are highly expressed *in planta* during infection of *A. australis* roots and leaves by *P. agathidicida* (Guo *et al*., 2020). PaRXLR24 triggers strong cell death in *N. benthamiana*, while PaRXLR40 suppresses immunity triggered by PaRXLR24 and other RXLR proteins in this plant species, indicating that its immune suppression activity is broad and not limited to a single elicitor (Guo *et al*., 2020). Despite this, the mechanisms by which these RXLRs influence disease outcomes - either by promoting virulence or, in some cases, triggering resistance - remain largely uncharacterized in *P. agathidicida*.

Here, we explore the functional role of the *P. agathidicida* RXLR effector PaRXLR40 and its potential to modulate host responses during infection. Based on its broad ability to suppress cell death in *N. benthamiana*, we hypothesized that PaRXLR40 acts as an immune suppressor that promotes pathogen colonization by interfering with host defence processes. To investigate this, we employed comparative approaches in both *N. benthamiana* and *A. australis*, aiming to uncover the molecular functions of PaRXLR40 and its contribution to host-pathogen interactions. We further hypothesized that PaRXLR40 may enhance virulence by targeting specific host proteins involved in immune regulation. To explore this possibility, we sought to identify candidate plant interactors and assess their potential role in modulating host responses. Additionally, we used RNA interference (RNAi) to silence *PaRXLR40* during infection and evaluate whether its silencing impairs *P. agathidicida* colonization. This study aims to provide insights into how individual RXLR effectors shape host susceptibility and disease progression.

## 2 Material and Methods

### 2.1 Microorganisms and plants

*P. agathidicida* strain 3770 (International Collection of Microorganisms (ICMP) 170237; Cox *et al*. (2022)) and *Phytophthora infestans* strain 88069 were used in this study. Wild-type *N. benthamiana* was grown in individual pots at 22°C with a 12 hour (h)/12 h light/dark cycle. *A. australis* plant material was derived from seeds originally sourced from Waipoua forest, NZ and grown at ambient temperature in a greenhouse then transferred to a shade house.

### 2.2 Yeast-two-hybrid assays

Yeast two-hybrid (Y2H) assays were conducted using the Invitrogen ProQuest system, as described by McLellan *et al*. (2021). Bait fusions were created via Gateway cloning and transformed into *Saccharomyces cerevisiae* MaV203 cells, where they were screened against a potato cDNA prey library (McLellan *et al*., 2013). Positive interactions were identified by growth on selective media lacking histidine or uracil and through β-galactosidase activity, followed by sequencing of interacting clones. Additional pairwise interaction tests were performed using wild-type and/or mutant bait/prey constructs. Primer sequences used for cloning are listed in Supp. Table S1.

### 2.3 *Agrobacterium tumefaciens*-mediated transient transformation assays

*Agrobacterium tumefaciens*-mediated transient transformation assays (ATTAs) were performed as described previously (Guo *et al*., 2020). In brief, ATTA expression vectors carrying the gene of interest were transformed into *A. tumefaciens* GV3101 (Holsters *et al*., 1980). Overnight cultures of *A. tumefaciens* transformants were suspended in infiltration buffer (10 mM MgCl_2_, 10 mM MES-KOH, pH 5.6) and infiltrated into the abaxial side of 4-week-old *N. benthamiana* leaves. For suppression, coimmunoprecipitation and total protein extraction assays, *A. tumefaciens* cultures with an OD_600_ of 0.5 were used, while for virulence assays, an OD_600_ of 0.1 was used, and for confocal microscopy experiments, an OD_600_ of 0.05 was used. For suppression assays, *A. tumefaciens* cultures carrying cell death elicitor ATTA expression vectors were infiltrated 24 h after the infiltration (hai) of *A. tumefaciens* cultures carrying cell death suppressor ATTA expression vectors. Primer sequences used for cloning are listed in Supp. Table S1.

### 2.4 Pathogenicity assays

*P. agathidicida* strain 3770 was subcultured onto cornmeal agar containing PARP (10 µg/ml (w/v) pimaricin, 250 µg/ml (w/v) ampicillin, 10 µg/ml (w/v) rifampicin, 100 µg/ml (w/v) pentachloronitrobenzene (PCNB)) (Morita & Tojo, 2007) as selective agents and grown at 22°C in the dark for 6 days. Then, 0.5 cm mycelium plugs were cut from the leading edge of actively growing cultures and inoculated culture-side down on the abaxial side of *N. benthamiana* or *A. australis* detached leaves. Inoculated leaves were kept in sealed plastic containers, lined with moist paper towels, to maintain humidity, and were incubated at 22°C with 12 h/12 h light/dark cycle.

For the colonization of *N. benthamiana* leaves and roots with *P. agathidicida*, *N. benthamiana* seeds were germinated in 24-well plates on sterile nappy liners soaked with Hoagland’s solution (Sigma-Aldrich) and incubated under a 12 h/12 h light/dark cycle at 22°C. After four weeks, *P. agathidicida* mycelial plugs were placed on the abaxial side of leaves or on roots, and seedlings were incubated in sealed petri dishes under the same conditions. Samples were collected at 6, 24, 48, and 72 hpi, matching timepoints used in a previous *P. agathidicida* gene expression study on *A. australis* (Cox et al., 2022). For each timepoint, three biological replicates were harvested, each consisting of pooled tissue from eight seedlings in contact with pathogen mycelium, which were rinsed and snap-frozen for downstream analyses.

*P. infestans* strain 88069 was maintained on rye agar plates at 19°C for two weeks. To harvest sporangia, each plate was flooded with 5 ml of sterile water and gently scraped with a glass rod. The suspension was transferred to a tube, sporangia were counted using a haemocytometer, and the concentration adjusted to 10^5^ cells/ml. Subsequently, 10 μl droplets of the suspension were applied to the abaxial side of *N. benthamiana* leaves, which were placed on damp tissue inside sealed containers to maintain humidity and incubated at room temperature.

### 2.5 Confocal microscopy

*A. tumefaciens* carrying a green fluorescent protein (GFP)-PaRXLR40 or red fluorescent protein (RFP)-StARIA expression construct was infiltrated into *N. benthamiana* leaves, as described above. Cells expressing these fluorescent protein fusions were observed using a Zeiss 710 confocal microscope at 2 days after infiltration (dai). GFP was excited at 488 nm, with emissions detected between 500 nm and 530 nm, while mRFP was excited at 561 nm, with emissions detected between 600 nm and 630 nm. On co-expression, GFP and RFP were imaged sequentially to minimize spectral cross-talk. Subsequent image processing for figure generation was conducted with the ImageJ software (Schindelin *et al*., 2012) and Adobe Illustrator.

### 2.5 Immunoprecipitation of tagged proteins from *Nicotiana benthamiana*

*A. tumefaciens* containing either the GFP-PaRXLR40 or RFP-StARIA fusion protein construct was infiltrated into *N. benthamiana* leaves as described above. Samples were collected 48 hai and total protein extracted as previously described (Guo *et al*., 2020). Protein fusions were immunoprecipitated using GFP-Trap-M magnetic beads (Chromotek), according to manufacturer’s instructions.

### 2.7 Western blotting

Total protein was extracted and subjected to western blot analysis as previously described (Guo *et al*., 2020). Proteins were separated by Sodium Dodecyl Sulphate-Polyacrylamide Gel Electrophoresis (SDS-PAGE) using 12% bis-tris-acrylamide separating gels with 5% stacking gels and transferred to polyvinylidene fluoride (PVDF) membranes. Detection was carried out using primary antibodies: PerCP-conjugated anti-GFP (mouse monoclonal, Santa Cruz sc-9996, 1:2000) and anti-RFP (rat monoclonal, Chromotek 5F8-150, 1:4000). The corresponding secondary antibodies were from LI-COR: IRDye® 800CW Goat anti-Mouse IgG (926-32210, 1:5000) and IRDye® 680LT Goat anti-Rat IgG (H+L) (926-68029, 1:5000). Protein bands were visualized using a LI-COR Odyssey CLX.

### 2.8 Endogenous application of ABA in *Nicotiana benthamiana* and *Agathis australis*

To modulate ABA levels in plant tissue, detached leaves of *N. benthamiana* and *A. australis* were treated with exogenous ABA. A 100 μM solution was prepared by diluting ABA (Sigma-Aldrich) in sterile distilled water containing 0.2% (v/v) ethanol. The solution was sprayed evenly onto *N. benthamiana* or *A. australis* leaves. Following ABA treatment, plants were maintained under high-humidity conditions and inoculated with *P. agathidicida* as described above. Control plants were sprayed with 0.2% (v/v) ethanol in water without ABA. Photos and infrared images were taken at 4 days post inoculation (dpi) and lesion areas were measured using ImageJ software (Schindelin *et al*., 2012).

### 2.9 RNA extraction and quantitative reverse transcription PCR

*N. benthamiana* leaves were infiltrated with *A. tumefaciens* carrying either an GFP-StARIA or free GFP expression vector, as described above. Leaf tissue samples were harvested 24 hai, and total RNA was extracted using a RNeasy Plant Mini Kit (Qiagen). RNA concentration and purity were assessed using a NanoDrop spectrophotometer (NanoDrop Technologies Inc.). For cDNA synthesis, 1 μg of total RNA per sample was used with the QuantiTect Reverse Transcription Kit (Qiagen). Quantitative polymerase chain reaction (qPCR) was carried out using the SensiFAST SYBR No-ROX mix (Meridian Bioscience) on a LightCycler 480 III (Roche). Relative expression levels of *N. benthamiana* genes were calculated using the 2^−ΔΔCt^ method (Livak & Schmittgen, 2001), with normalization to the reference gene *NbActin* (Sainsbury & Lomonossoff, 2008). Expression values were calculated relative to the GFP control, which was set to 1. Data are presented as means ± standard error from three independent biological replicates. Statistical significance was assessed using Student’s *t*-test. Primer sequences used for qPCR are listed in Supp. Table S1.

### 2.10 Synthesis of dsRNA constructs and detached leaf assay with dsRNA

Sequences of *GFP* and *PaRXLR40* targeted for gene silencing using double-stranded RNA (dsRNA) (714 bp for *GFP* and 476 bp for *PaRXLR40*) were amplified from plasmid pB7WGF2 (Karimi *et al*., 2002) and *P. agathidicida* cDNA, respectively, using primers containing T7 promoter sequences at the 5’ end (Supp. Table S1). dsRNA was synthesized using a MEGAScript^TM^ RNAi kit (Invitrogen).

dsRNA^GFP^ or dsRNA^PaRXLR40^ were diluted to 20 ng/µL in TE buffer (10 µM Tris-HCl and 1 µM EDTA, pH 8.0) plus 0.1% (v/v) of Tween 20 and were either sprayed into the abaxial side of detached *N. benthamiana* leaves or infiltrated into detached *A. australis* leaves (Video S1). Leaves were kept in sealed plastic containers, lined with moist paper towels, and incubated at 22°C in the dark. Twenty-four h later, the right and left side of *N. benthamiana* leaves and each *A. australis* leaf were inoculated with *P. agathidicida* as described above and incubated at 22°C with 12 h/12 h light/dark cycle. For *N. benthamiana*, one biological replicate was defined as a detached leaf inoculated on both sides. Three biological replicates were used for qPCR analysis, in which tissue from both inoculation sites on each leaf was pooled for RNA extraction. Eight biological replicates were used for lesion measurement, with lesion size assessed separately for each side of the leaf and average per replicate. For *A. australis*, each leaf represented one biological replicate; five leaves were used for qPCR and 17 for lesion measurement in experiment 1, while experiment 2 included three biological replicates for qPCR and eight for lesion measurement.

For *N. benthamiana*, samples were collected at 24 hpi for gene expression analysis, while lesion area and pictures were taken at 72 hpi. For *A. australis*, samples were collected at 48 hpi for gene expression analysis, while lesion area and pictures were taken at 5 dpi. Total RNA was extracted from infected *A. australis* tissue using a combined CTAB and column-based method. Briefly, tissue was ground in liquid nitrogen and incubated in pre-heated CTAB extraction buffer (3% CTAB, 3% PVP-40, 100 mM Tris-HCl pH 8.0, 25 mM EDTA, 2 M NaCl, 0.5 g/L spermidine, and 2% β-mercaptoethanol) at 65 °C for 10 min. The lysate was then extracted twice with chloroform:isoamyl alcohol (24:1) and the aqueous phase recovered. RNA was then purified using the AllPrep® Fungal DNA/RNA/Protein Kit (QIAGEN), following the manufacturer’s protocol. Expression of *PaRXLR40* was calculated by qPCR using the 2^−ΔCt^ method (Livak & Schmittgen, 2001), with normalization to the reference genes *PaActin* (KNV87_002159-T1) and *Paβ-tubulin* (KNV87_012438-T1; Cox *et al*. (2022)). This approach yields normalized relative quantities (NRQ), representing the abundance of *PaRXLR40* transcripts relative to internal controls within each sample. Data are presented as means ± standard error and results were statistically analysed using Student’s *t* test. Primer sequences used for qPCR are listed in Supp. Table S1.

## 3 Results

### 3.1 PaRXLR40 is a broad cell death suppressor and requires its C-terminal region for activity

Using ATTAs in *N. benthamiana*, we previously determined that PaRXLR40 suppresses cell death triggered by the *P. agathidicida* effector, PaRXLR24, as well as by the *P. infestans* RXLR effector Avr3a in the presence of its cognate potato immune receptor protein R3a (Guo *et al*., 2020). Given these results, we tested whether PaRXLR40 could also suppress cell death triggered by a characterized glycoside hydrolase family 12 (GH12) effector from *P. agathidicida*, PaXEG1, which is a homologue of a cell death-eliciting effector from *Phytophthora sojae* (Bradley, 2022). Here, suppression of PaRXLR24- and Avr3a/R3a-elcitied cell death by PaRXLR40 were used as controls (Fig. 1a and 1b). Interestingly, PaRXLR40 also suppressed cell death triggered by PaXEG1, suggesting that PaRXLR40 can suppress cell death triggered by perception of both intracellular and extracellular (apoplastic) effector proteins (Fig. 1c).

**Figure 1.**
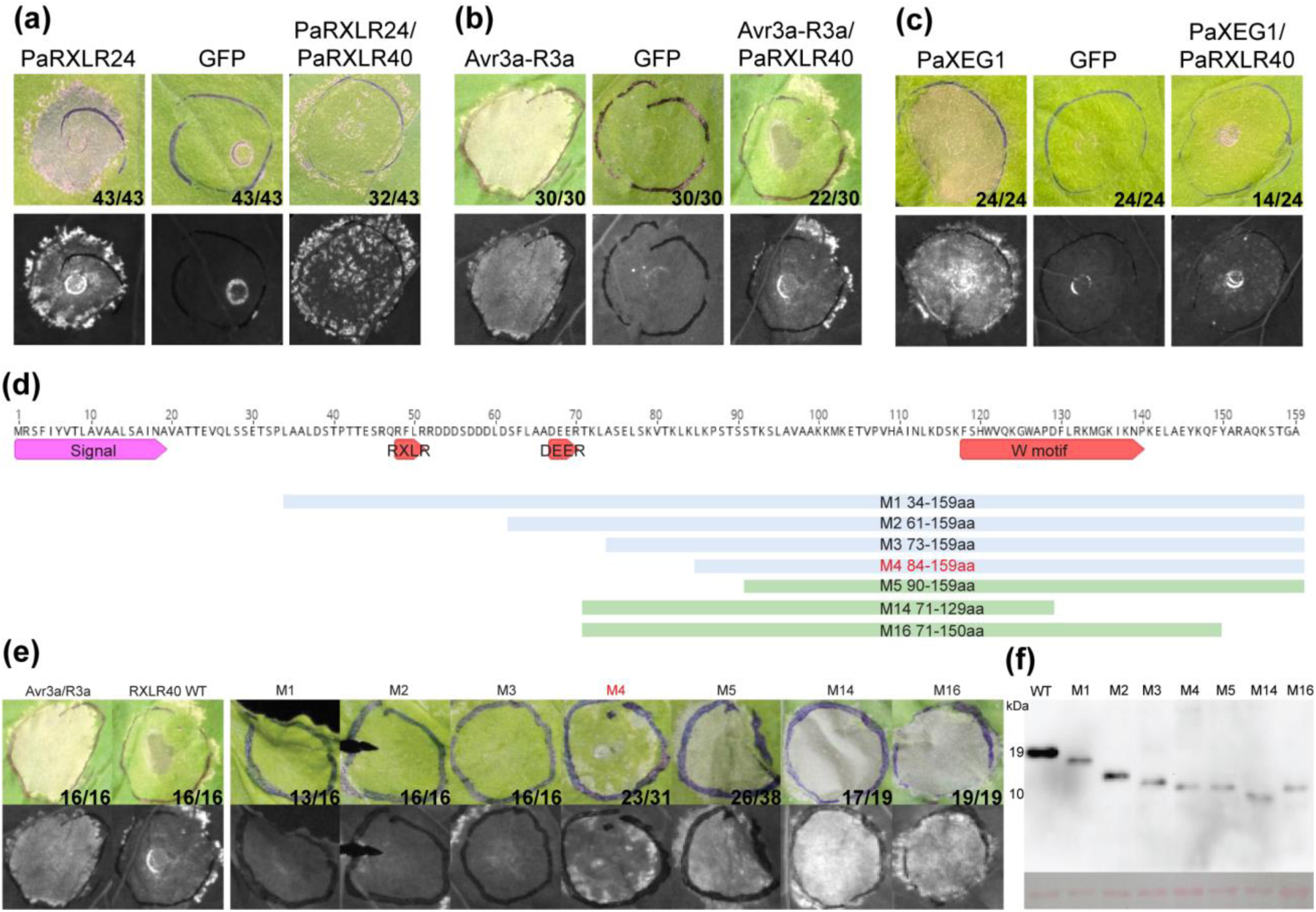
PaRXLR40 of *Phytophthora agathidicida* suppresses both intracellular and extracellular cell death elicitors. Suppression of (a) PaRXLR24-triggered cell death, (b) Avr3a/R3a-triggered cell death and (c) PaXEG1-triggered cell death by PaRXLR40 in *Nicotiana benthamiana*. *Agrobacterium tumefaciens* carrying cell death elicitor expression vectors were infiltrated 1 day after infiltration (dai) of *A. tumefaciens* carrying the PaRXLR40 expression vector. Photographs with visible (top) and UV (bottom) light were taken 6 days after the second infiltration step. Representative images are shown from four independent experiments. Numbers on the bottom right-hand side represent the number of times the response was observed (left) out of the number of times the agroinfiltration was performed (right). Experiments in (a) and (b) were repeated according to Guo *et al*. (2020). (d) Schematic diagram of PaRXLR40 truncation mutations. The pink arrow shows the predicted signal peptide. The red arrows indicate the RXLR, EER and W motifs of PaRXLR40. The blue lines show truncated mutants of PaRXLR40 (M1-4) that could still suppress Avr3a/R3a-triggered cell death. The green lines show truncated mutants of PaRXLR40 (M5, M14 and M16) that could not suppress Avr3a/R3a-triggered cell death. (e) *A. tumefaciens* carrying wild-type (WT) or truncated PaRXLR40 mutant (M) expression vectors were co-infiltrated with *A. tumefaciens* carrying cell death elicitor Avr3a/R3a expression constructs to determine the regions of PaRXLR40 required for suppression of cell death. *A. tumefaciens* carrying Avr3a/R3a expression vectors were infiltrated 1 dai of *A. tumefaciens* carrying WT or truncated PaRXLR40 expression vectors. Photographs with visible (top) and UV (bottom) light were taken 6 days after the second infiltration step. From left to right: Avr3a/R3a-triggered cell death; PaRXLR40-mediated suppression of Avr3a/R3a-triggered cell death (positive control); PaRXLR40 M1-M4-mediated suppression of Avr3a/R3a-triggered cell death; failed suppression of Avr3a/R3a-triggered cell death by PaRXLR40 M5, M14 and M16. The experiment was repeated at least three times with consistent results. Numbers on the bottom right-hand side represent the number of times the response was observed (left) out of the number of times the agroinfiltration was performed (right). (f) Protein immunoblots of total proteins extracted from *N. benthamiana* leaves collected 3 dai confirmed the presence of WT and truncated PaRXLR40 constructs (M).

To determine the regions of PaRXLR40 required for the suppression of Avr3a/R3a-triggered cell death, truncated FLAG-tagged versions of PaRXLR40 were generated (Fig. 1d) and tested using ATTAs in *N. benthamiana*. The results showed that PaRXLR40 mutants (M)1 (missing the first 33 aa), M2 (missing the first 60 aa, including the RXLR motif), M3 (missing the first 72 aa, including the RXLR and EER motifs) and M4 (missing the first 83 aa, including the RXLR and EER motifs), provided similar levels of cell death suppression to the wild-type effector, whilst M5, which is missing the first 90 aa (including the RXLR and DEER motifs) failed to suppress cell death (Fig. 1e). Furthermore, M14, which is missing the first 70 aa and last 30 aa (including the RXLR and EER motif, as well as part of the WY domain), and M16, which is missing the first 70 aa and last 9 aa (including the RXLR and EER motif), also failed to suppress Avr3a/R3a triggered cell death (Fig. 1e). Given that all mutants could be detected by western blotting (Fig. 1f), these results suggest that the RXLR and EER motifs of PaRXLR40 are not required for suppression activity, but that the C-terminal region (84-159 aa) of PaRXLR40, which contains the WY domain, is required for suppression activity.

### 3.2 PaRXLR40 enhances *Phytophthora* infection in *Nicotiana benthamiana*

To examine *PaRXLR40* expression over the course of host infection, we analysed the transcriptomic dataset from *P. agathidicida*-infected *A. australis* (Cox *et al*., 2022) and the expression of *PaRXLR40* across different infection time points in *N. benthamiana*. In *A. australis*, *PaRXLR40* was expressed in both roots and leaves at all time points, with no statistically significant differences in expression levels across samples (Fig. S1a). In contrast, in *N. benthamiana*, *PaRXLR40* was more strongly expressed in leaves than in roots (Fig. S1b). Leaf expression peaked at 24 h post inoculation (hpi) and was significantly higher than root expression at the same time point. These results suggest that *PaRXLR40* is expressed throughout infection in both hosts but shows host- and tissue-specific regulation.

Next, we questioned whether PaRXLR40 is a virulence factor that facilitates *P. agathidicida* colonization of *N. benthamiana* leaves. Transient expression of GFP-PaRXLR40 in *N. benthamiana* leaves, followed by pathogen inoculation, resulted in significantly larger *P. agathidicida* lesions compared with GFP expression, while expression of GFP-PaRXLR40 M14, the loss of suppression mutant, had no effect on *P. agathidicida* infection when compared with the GFP control (Fig. 2). Similar results were observed when these genes were overexpressed in *N. benthamiana* and inoculated with *P. infestans* (Fig. S2), suggesting that PaRXLR40 might target a conserved pathway or molecule in *N. benthamiana*.

**Figure 2.**
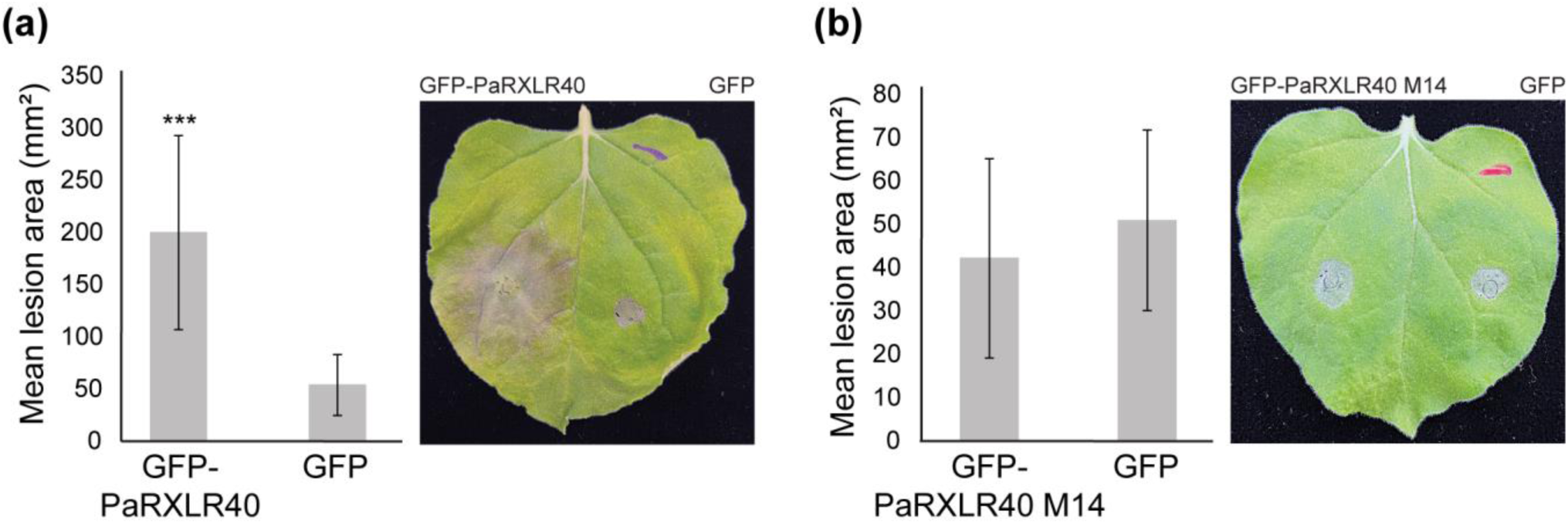
PaRXLR40 enhances *Phytophthora agathidicida* infection in *Nicotiana benthamiana*. Overexpression of (a) GFP-PaRXLR40 in *N. benthamiana* enhances *P. agathidicida* leaf colonization, while overexpression of (b) PaRXLR40 mutant 14 (M14) shows similar levels of infection observed for the GFP control. *Agrobacterium tumefaciens* carrying expression vectors for each protein were infiltrated in opposing leaf segments of *N*. *benthamiana*. Leaves were then inoculated with *P*. *agathidicida* at 1 day after infiltration (dai) with *A. tumefaciens*. Photos and measurements of lesion area (mm^2^) were taken at 4 days after inoculation. Means and standard errors were calculated from four biological replicates. ***, *P*<0.001 using Student’s *t*-test.

### 3.3 Silencing of PaRXLR40 reduces Phytophthora agathidicida infection in Nicotiana benthamiana and Agathis australis

Since PaRXLR40 enhances *P. agathidicida* infection in *N. benthamiana*, we investigated whether silencing this effector using a dsRNA-based RNAi approach would reduce *P. agathidicida* virulence. In *N. benthamiana*, dsRNA targeting *PaRXLR40* (dsRNA^PaRXLR40^) was applied by foliar spray prior to pathogen inoculation (Kalyandurg *et al*., 2021). To confirm silencing efficiency, *PaRXLR40* transcript levels were quantified by qPCR in infected leaf tissue. *PaRXLR40* transcript levels were significantly reduced in dsRNA^PaRXLR40^-treated leaves compared to the dsRNA^GFP^ control (Fig. 3a). To determine whether the reduced *PaRXLR40* expression affected pathogen virulence, lesion areas were measured in *N. benthamiana* leaves treated with dsRNA^PaRXLR40^ or control dsRNA^GFP^. dsRNA^PaRXLR40^ significantly reduced lesion size compared to dsRNA^GFP^ (Fig. 3b,c).

**Figure 3.**
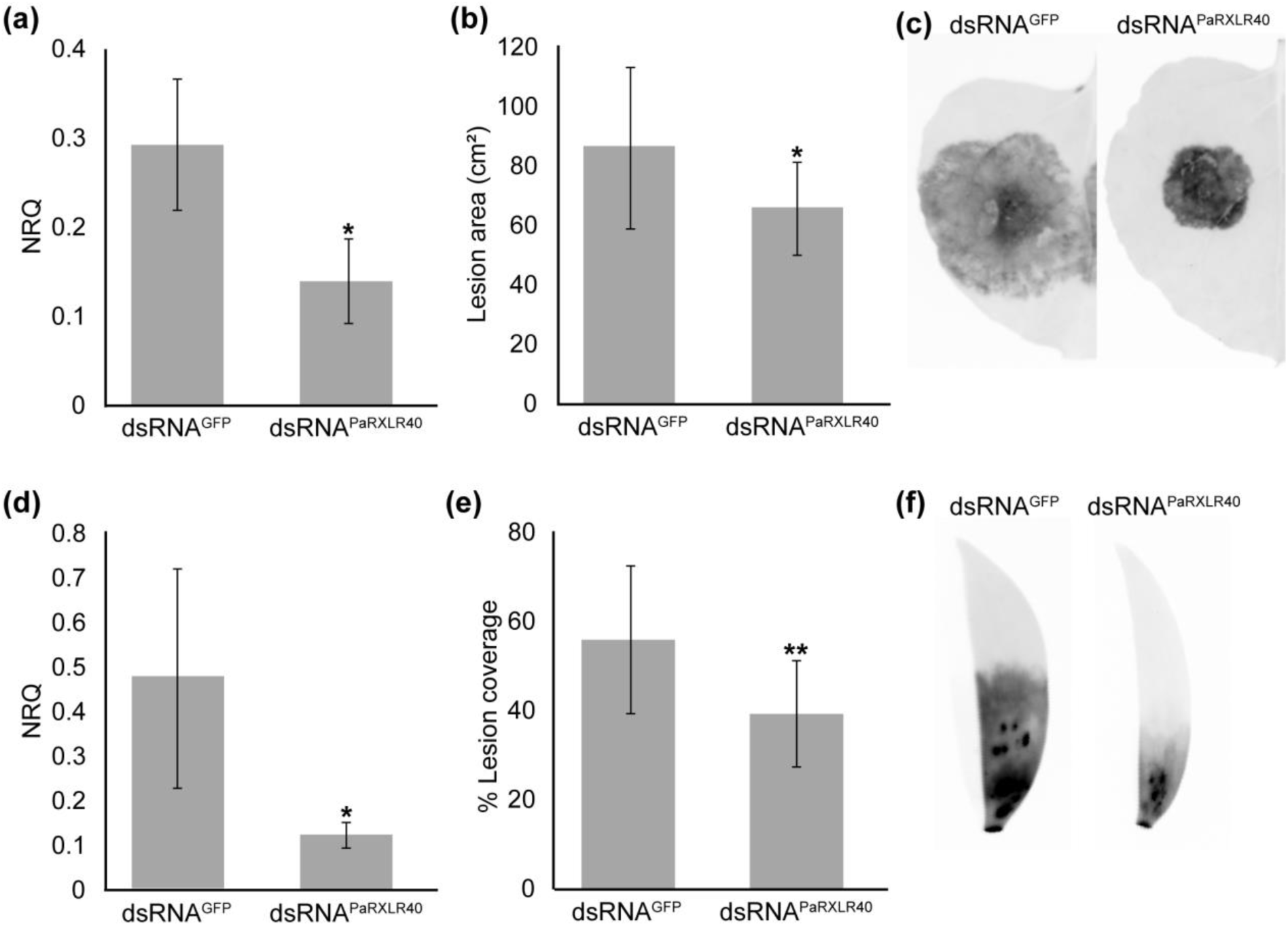
Silencing of *PaRXLR40* reduces *Phytophthora agathidicida* lesion size in *Nicotiana benthamiana* and *Agathis australis*. (a, d) Relative expression of *PaRXLR40* in *P. agathidicida*-infected leaves of (a) *N. benthamiana* at 24 h post inoculation (hpi) and (d) *A. australis* at 48 hpi, following prior treatment with dsRNA^PaRXLR40^ or dsRNA^GFP^. Here, dsRNA leaf treatments were carried out 24 h before pathogen inoculation. Normalized relative quantity (NRQ) values represent normalized relative quantification of *PaRXLR40* transcript levels, calculated using *PaActin* and *Paβ-tubulin* as reference genes. Means and standard errors were calculated from three biological replicates in *N. benthamiana* and five in *A. australis*. (b, e) Quantification of disease symptoms in (b) *N. benthamiana* as lesion area (cm²) and (e) *A. australis* as percentage (%) lesion coverage. Lesion coverage was quantified as the percentage of the leaf area affected by disease, calculated as the ratio between lesion length and total leaf length for each sample. Means and standard errors were calculated from eight biological replicates in *N. benthamiana* and 17 in *A. australis*. *, *P*<0.05; **, *P*<0.01 using Student’s *t*-test. (c, f) Representative infrared images of disease symptoms at 72 hpi in (c) *N. benthamiana* and at 5 days post inoculation in (f) *A. australis*.

To test whether this effect was conserved in *A. australis*, detached leaves were infiltrated with dsRNA^PaRXLR40^ or dsRNA^GFP^ treatments prior to pathogen inoculation. Here, leaves treated with dsRNA^PaRXLR40^ showed significantly reduced *PaRXLR40* transcript levels compared to the dsRNA^GFP^, confirming effective silencing (Fig. 3d). This reduction correlated with smaller lesion coverage, indicating decreased pathogen colonization (Fig. 3e,f). This same experiment was repeated a second time and similar results were observed (Fig. S3). These results demonstrate that *PaRXLR40* contributes to *P. agathidicida* virulence in *N. benthamiana* and *A. australis* and highlights its potential as a target for RNAi-based disease control strategies.

### 3.4 PaRXLR40 interacts with StARIA

To investigate the role that PaRXLR40 plays in promoting *P. agathidicida* virulence *in planta*, a yeast-2-hybrid (Y2H) screen was conducted using a GAL4 DNA-binding domain fusion with PaRXLR40 as the bait. The screen employed a cDNA library derived from *Solanum tuberosum* infected with *Phytophthora infestans* (Bos *et al*., 2010; McLellan *et al*., 2022) and reached a coverage of 4.24 × 10⁶ yeast co-transformants. From the selection plates, eight positive yeast colonies expressing GAL4 activation domain (prey) fusions were recovered, all of which corresponded to ARIA (arm repeat protein interacting with ABF2), a BTB/POZ domain-containing protein, hereafter referred to as StARIA (Table S2). In *Arabidopsis*, AtARIA has been described as a positive regulator of ABA responses (Kim *et al*., 2004) and has 75.4% pairwise amino acid identity with StARIA (Fig. S4). We also identified the orthologue of StARIA in *N. benthamiana* (NbARIA – Niben101Scf02021g02009.1), with 93.7% pairwise identity between them (Fig. S4). Next, we analysed the expression of *NbARIA* during *P. agathidicida* infection in leaves and roots of *N. benthamiana*. In leaves, *NbARIA* expression was highest at 6 hpi, with reduced levels at 24 and 48 hpi (Fig. S5). In contrast, expression levels in roots remained relatively stable across all time points and were consistently lower than those observed in leaf tissue (Fig. S5). This suggests a possible tissue-specific pattern of *NbARIA* expression at an early stage of *P. agathidicida* infection.

To validate the interaction between PaRXLR40 and StARIA, a full-length *StARIA* prey construct was tested in pairwise Y2H assays with bait constructs encoding PaRXLR40, PaRXLR40 M14 (loss of suppression mutant), PaRXLR24 and an empty bait vector. While all yeast transformants were able to grow on non-selective control medium, only the combination of PaRXLR40 and StARIA supported growth on selective medium and activation of the β-galactosidase reporter gene (Fig. 4a and Fig. S6), indicating a specific interaction. To assess whether this interaction also occurs *in planta*, co-immunoprecipitation was performed in *N. benthamiana*. RFP-tagged StARIA was expressed alone or co-expressed with either GFP-PaRXLR40 or PaRXLR40 M14, and immunoprecipitation was conducted using GFP-Trap magnetic beads. As shown in Fig. 4b, all proteins were present in the respective input samples. RFP-StARIA was co-immunoprecipitated in the presence of GFP-PaRXLR40, but not when expressed with PaRXLR24; however, it was also co-immunoprecipitated in the presence of PaRXLR40 M14 (Fig. 4b), which suggests that the interaction region of PaRXLR40 with StARIA might be different from the region required for cell death suppression.

**Figure 4.**
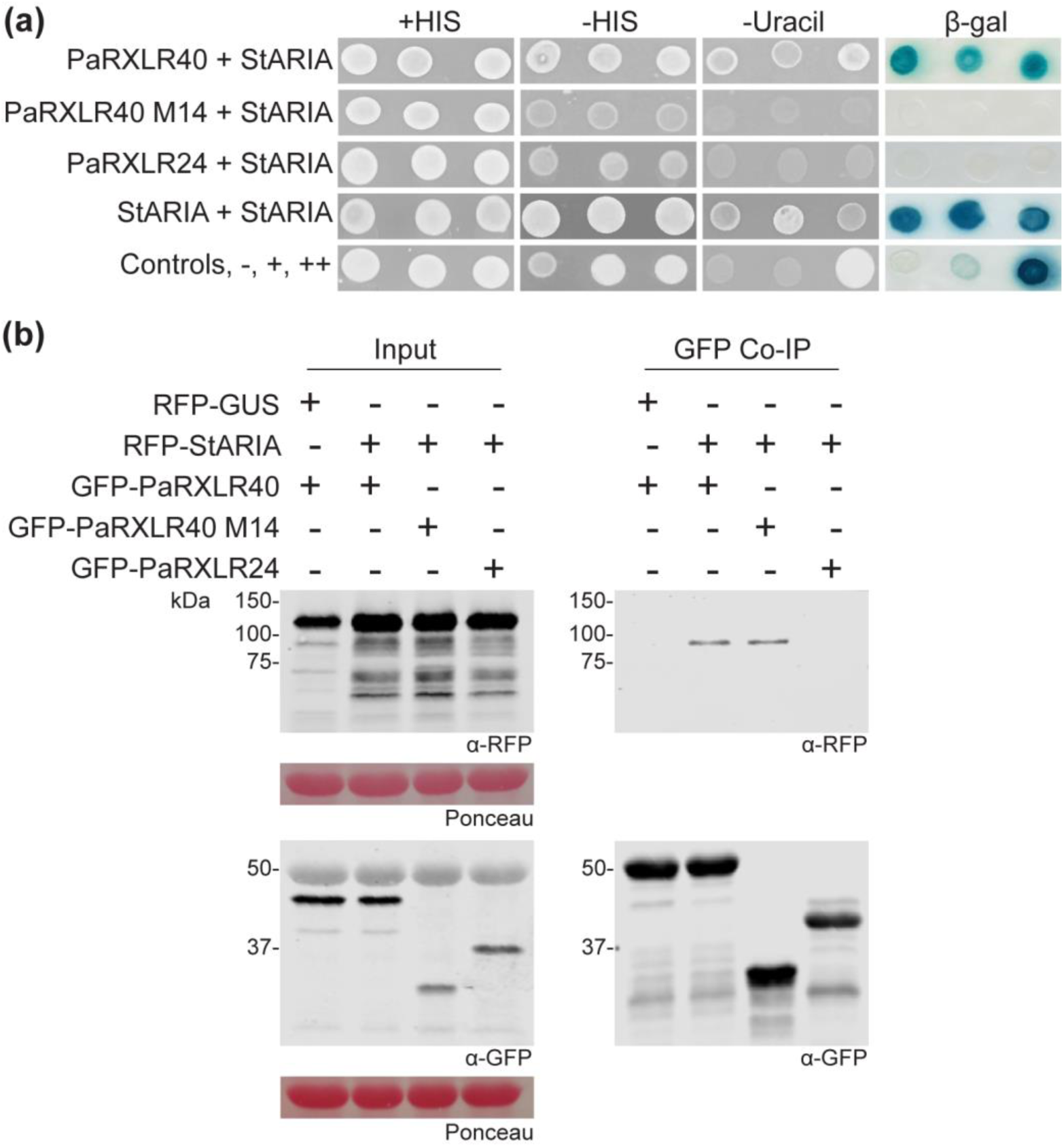
PaRXLR40 interacts with StARIA *in vitro* and *in planta*. (a) Yeast co-expressing PaRXLR40 and StARIA, or StARIA and StARIA (triplicate samples) were able to grow on selective medium lacking histidine (-HIS) and uracil (-Uracil) and exhibited β-galactosidase (β-gal) activity, indicating a positive interaction. In contrast, yeast co-expressing PaRXLR40 M14 or PaRXLR24 with StARIA failed to grow on −HIS and −Uracil medium and showed no reporter activity. Growth on +HIS medium confirmed that all yeast strains were viable under non-selective conditions. Yeast controls were: −, no interaction; +, weak interaction; ++, strong interaction. (b) Co-immunoprecipitation (Co-IP) from *N. benthamiana* leaf tissue confirmed that RFP-tagged StARIA interacted with GFP-tagged PaRXLR40, but not with GFP-PaRXLR24, however it also interacted with GFP-PaRXLR40 M14. RFP-GUS (β-glucuronidase) was included as a non-interacting negative control. Protein extracts from agroinfiltrated leaves were subjected to immunoprecipitation using GFP-Trap beads. Construct expression in leaves is indicated by “+”, and molecular weight markers (in kDa) are shown. Protein loading was verified by Ponceau S staining.

We also determined that StARIA interacts with itself *in vitro*, likely forming a dimer (Fig. 4a), something that has been shown for BTB-containing proteins in other eukaryotes (Stogios & Privé, 2004). A mutation at a conserved aspartate residue (D35) in PLZF, a BTB/POZ domain-containing protein, was previously shown to reduce dimerization efficiency (Melnick *et al*., 2000). Alignment of several BTB/POZ domains with that of StARIA revealed a conserved aspartate residue (D542). To assess the role of this residue in StARIA, D542 was substituted with alanine (StARIA^D542A^), and the impact on dimerization evaluated using a Y2H assay. Results showed that yeast co-expressing StARIA^D542A^ and wild-type StARIA failed to grow on media lacking histidine or uracil and showed no β-galactosidase activity. This indicated that substitution of D542 disrupts StARIA dimerization (Fig. S6). Additionally, PaRXLR40 did not interact with StARIA^D542A^, suggesting that dimerization may be necessary for interaction with the effector (Fig. S6).

To determine the subcellular localization of PaRXLR40 and its host interactor StARIA *in planta*, GFP-RXLR40, GFP-StARIA or RFP-StARIA were transiently expressed in *N. benthamiana* leaves. Confocal microscopy revealed that both proteins localize predominantly to the nucleus, as confirmed by co-localization with the nuclear marker CFP-H2B (cyan fluorescent protein fused to histone H2B) (Fig. 5 and Fig. S8). Notably, GFP-RXLR40 also localised in the nucleolus, with the same pattern for PaRXLR40 M14 (Fig. S7 and Fig. S8), whereas GFP-StARIA appeared to be excluded from this region. Co-expression of GFP-RXLR40 and RFP-StARIA did not alter the localization patterns of either protein (Fig. 5 and Fig. S8), indicating that their interaction does not occur in the nucleolus.

**Figure 5.**
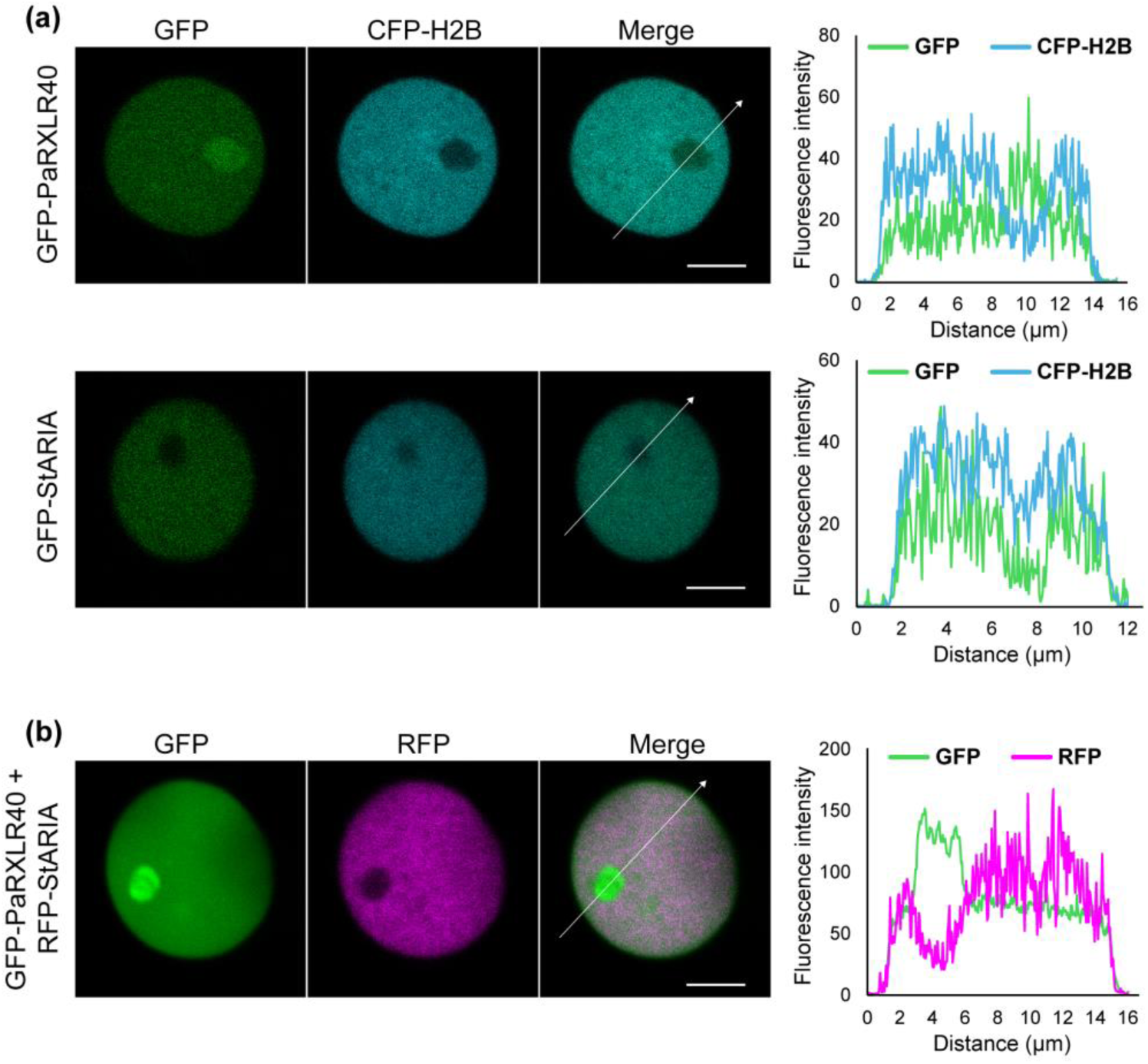
PaRXLR40 and StARIA co-localize to the nucleus of *Nicotiana benthamiana*. (a) Representative confocal images of *N. benthamiana* epidermal cells transiently expressing GFP-PaRXLR40 or GFP-StARIA, together with CFP-H2B (nuclear marker; CFP fused to histone H2B). GFP-PaRXLR40 localizes to the nucleus and shows enrichment in the nucleolus, while GFP-StARIA localizes to the nucleoplasm but is excluded from the nucleolus. (b) Co-expression of GFP-PaRXLR40 and RFP-StARIA shows overlapping nuclear localization without altered distribution patterns. White arrows indicate the transects used to generate the fluorescence intensity plots shown to the right of each image set. The X-axis of each plot represents the distance (in µm) along the corresponding white arrow. Scale bars = 5 µm.

### 3.5 StARIA overexpression suppresses effector-triggered cell death and enhances *Phytophthora agathidicida* infection in *Nicotiana benthamiana*

Since PaRXLR40 is known to suppress effector-triggered cell death, we investigated whether its host target, StARIA, altered effector-triggered cell death. To test this, we evaluated whether overexpression of *StARIA* could suppress or enhance cell death induced by the previously tested cell death elicitors in *N. benthamiana*. Transient expression of *StARIA* reduced cell death triggered by PaRXLR24, Avr3a/R3a, and PaXEG1, when *A. tumefaciens* carrying expression vectors for these effectors were infiltrated 24 h after *A. tumefaciens* carrying the *StARIA* expression vector (Fig. 6). In contrast, strong cell death responses were observed when effectors were expressed alongside the GFP control. These results indicate that StARIA, like PaRXLR40, functions as a suppressor of effector-triggered cell death *in planta*.

**Figure 6.**
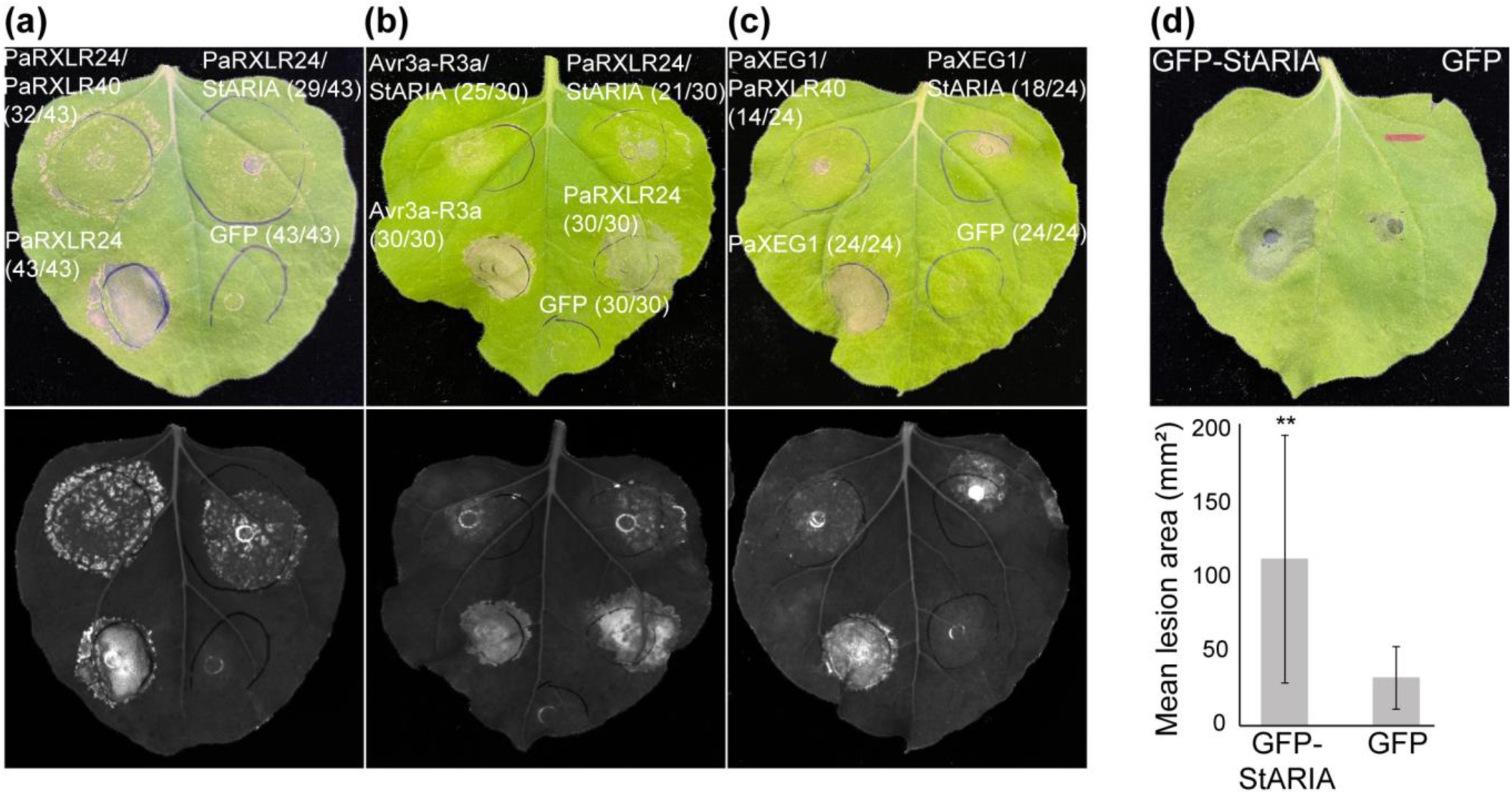
StARIA suppresses cell death and enhances *Phytophthora agathidicida* infection in *Nicotiana benthamiana*. Suppression of (a) PaRXLR24-triggered cell death, (b) Avr3a/R3a-triggered cell death and (c) PaXEG1-triggered cell death by StARIA in *N. benthamiana*. PaRXLR24/StARIA was included in (b) and PaXEG1/PaRXLR40 was included in (c) as suppression controls. *Agrobacterium tumefaciens* carrying expression vectors for cell death elicitors were infiltrated 1 day after infiltration (dai) of *A. tumefaciens* carrying an expression vector for StARIA. Photographs with visible (top) and UV (bottom) light were taken 6 dai of *A. tumefaciens* carrying the cell death elicitor expression vectors. Representative images are shown from four independent experiments. Numbers in parentheses in (a-c) indicate the number of times the response was observed (left) out of the number of times the agroinfiltration was performed (right). (d) Overexpression of GFP-StARIA in *N. benthamiana* enhances *P. agathidicida* leaf colonization. *A. tumefaciens* carrying expression vectors for each protein were infiltrated into opposing leaf segments of *N*. *benthamiana*. Leaves were then inoculated with *P*. *agathidicida* 1 dai with *A. tumefaciens*. Photos and measurements of lesion area (mm^2^) were taken at 4 days post inoculation. Means and standard errors were calculated from four biological replicates. **, *P*<0.01 using Student’s *t*-test.

To determine whether this suppression enhances the pathogen colonization of *N. benthamiana*, leaves expressing either GFP-StARIA or GFP were challenged with *P. agathidicida*. At 4 dpi, significantly larger lesions were observed in leaves expressing GFP-StARIA compared to the GFP control (Fig. 6d), indicating increased susceptibility. Together, these findings suggest that StARIA contributes to *P. agathidicida* virulence by suppressing host cell death responses.

Interestingly, mutation of the conserved aspartate residue within the BTB domain, predicted to disrupt dimerization, did not impair the ability of StARIA to suppress effector-triggered cell death or promote *P. agathidicida* infection in *N. benthamiana* (Fig. S9), suggesting that StARIA may function independently of BTB-mediated dimerization or employ alternative mechanisms. There is also the possibility that the endogenous wild-type version of ARIA from *N. benthamiana* could still dimerize with the mutant form, contributing to the observed suppression phenotype.

### 3.6 Abscisic acid enhances *P. agathidicida* infection

Since ARIA has been described as a positive regulator of ABA responses in *Arabidopsis* (Kim *et al*., 2004), we investigated the effect of ABA on *P. agathidicida* infection. In *N. benthamiana*, ABA treatment led to a significant increase in lesion area compared to mock-treated control plants (Fig. 7a), with visibly larger lesions at 4 dpi. Similarly, in *A. australis*, ABA-treated leaves developed significantly larger lesions than mock-treated controls at 4 dpi (Fig. 7b). These results suggest that exogenous ABA enhances *P. agathidicida* infection in both hosts. Next, we hypothesized that ABA may influence host susceptibility to pathogens through modulation of ABA signalling. To investigate this, we analysed the expression of some key ABA-related genes following StARIA overexpression in *N. benthamiana*. Transcript levels of 9-cis-epoxycarotenoid dioxygenase 3 (*NbNCED3*) and ABA-deficient 2 (*NbABA2*), both involved in ABA biosynthesis, were significantly upregulated compared to GFP controls (Fig. 7c,d). These findings suggest that StARIA may contribute to increased host susceptibility to pathogens by modulating ABA biosynthesis, supporting a potential role for StARIA in facilitating ABA-dependent suppression of plant immune responses.

**Figure 7.**
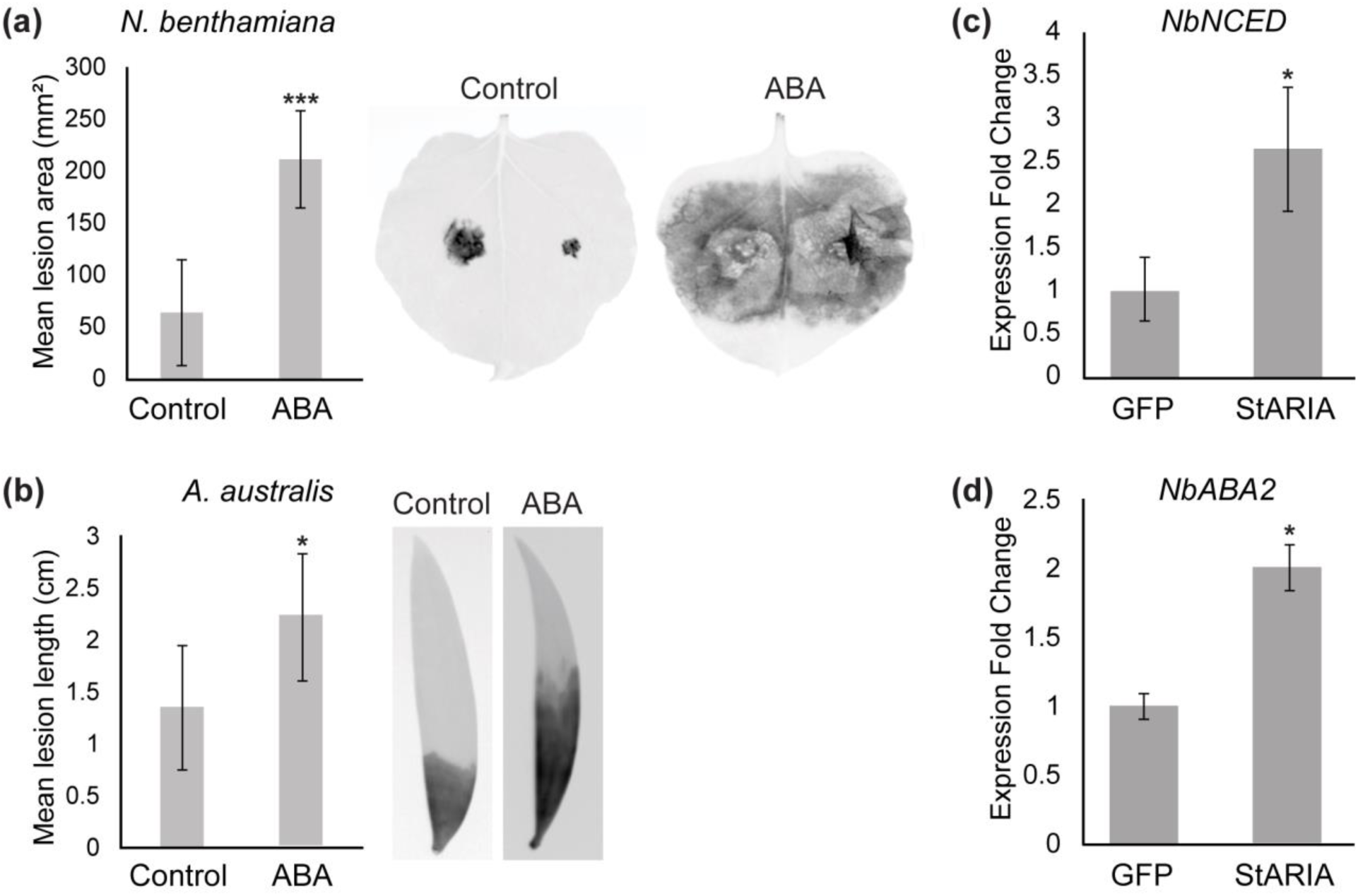
ABA promotes infection of host plants by *Phytophthora agathidicida* and StARIA upregulates ABA-related gene expression in *Nicotiana benthamiana*. (a) *N. benthamiana* and (b) *Agathis australis* detached leaves were treated with an abscisic acid (ABA) spray to a final concentration of 100 µM or a mock-treated control, followed by *P. agathidicida* inoculation. Graphs shows mean lesion area (mm^2^) in *N. benthamiana* (a) and mean lesion length (cm) in *A. australis* (b). Infrared photos and measurements of lesion area (mm^2^) were taken at 4 days post-inoculation. Means and standard errors were calculated from 35 detached leaves. *, *P*<0.05, ***, *P*<0.001 using Student’s *t*-test. (c) Expression of ABA-related genes, *NbNCED3* and *NbABA2*, in *N. benthamiana*. GFP-StARIA and GFP control were transiently expressed in *N. benthamiana* and leaves sampled 24 h after infiltration. Means and standard errors of normalised (*NbActin*) expression values were calculated from three biological replicates. *, *P*<0.05, using Student’s *t*-test.

### 3.7 StARIA interacts with a NAC transcription factor

To identify potential host interactors of StARIA, we performed another Y2H screen with StARIA as the bait. The screen reached a coverage of 2.2 × 10^5^ yeast co-transformants and seven positive yeast colonies were recovered from selection plates, corresponding to SOG1 (SUPPRESSOR OF GAMMA RESPONSE 1) hereafter referred to as StSOG1 (Fig. S10; Table S2). In *Arabidopsis*, AtSOG1 has been described as NAC-domain transcription factor that plays a central role in the DNA damage response (Yoshiyama *et al*., 2009) and has 57.5% pairwise amino acid identity with StSOG1 (Fig. S10). The orthologue of StSOG1 in *N. benthamiana* (NbSOG1 – Niben101Scf01627g02012.1; Fig. S10) was used to validate the interaction.

Here, a full-length *NbSOG1* prey construct was tested in pairwise Y2H assays with bait constructs encoding StARIA, PaRXLR40 and an empty vector. While all yeast transformants were able to grow on non-selective control medium, only the combination of NbSOG1 and StARIA supported growth on selective medium and activation of the β-galactosidase reporter gene (Fig. 8a and Fig. S6), indicating a specific interaction *in vitro*.

**Figure 8.**
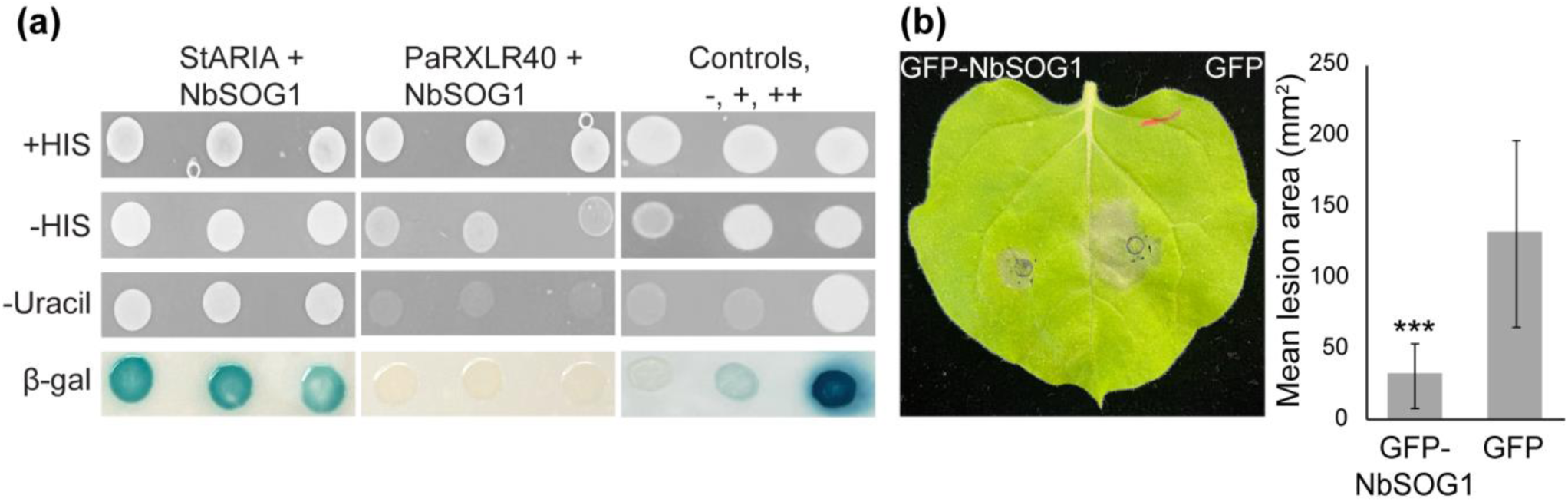
NbSOG1 interacts with StARIA and enhances resistance to *Phytophthora agathidicida* in *Nicotiana benthamiana*. (a) Yeast co-expressing StARIA and NbSOG1 were able to grow on selective medium lacking histidine (-HIS) and uracil (-Uracil) and exhibited β-galactosidase (β-gal) activity, indicating a positive interaction. In contrast, yeast co-expressing PaRXLR40 and NbSOG1 failed to grow on −HIS and −Uracil medium and showed no reporter activity. Growth on +HIS medium confirmed that all yeast strains were viable under non-selective conditions. Yeast controls were: −, no interaction; +, weak interaction; ++, strong interaction. (b) Overexpression of GFP-NbSOG1 in *N. benthamiana* reduces *P. agathidicida* leaf colonization. *Agrobacterium tumefaciens* carrying expression constructs for each protein were infiltrated into opposing leaf segments of *N*. *benthamiana*. Leaves were then inoculated with *P*. *agathidicida* at 1 days after infiltration with *A. tumefaciens*. Photos and measurements of lesion area (mm^2^) were taken 4 days post inoculation. Means and standard errors were calculated from four biological replicates. ***, *P*<0.001 using Student’s *t*-test.

Since AtSOG1 has been described as positively regulating immune responses against pathogens (Yoshiyama *et al*., 2020), we wanted to determine whether *NbSOG1* overexpression would enhance resistance against *P. agathidicida* in *N. benthamiana*. At 4 dpi, significantly smaller lesions were observed in leaves expressing GFP-NbSOG1 compared to the GFP control (Fig. 8b), indicating increased resistance.

## 4 Discussion

Effector proteins secreted by *Phytophthora* species play essential roles in overcoming host defences and promoting disease. Here, we investigated the *P. agathidicida* RXLR effector PaRXLR40, which appears to be unique to *P. agathidicida* and broadly conserved among its isolates, as well as being highly expressed in infected *N. benthamiana* and *A. australis* tissues. Although PaRXLR40 is expressed during infection of its gymnosperm host, *A. australis*, its virulence-promoting activity has not yet been directly tested in this system. Due to the cultural significance of *A. australis* and technical limitations that result from the absence of a sequenced genome and established molecular tools for *A. australis*, we employed the model angiosperm *N. benthamiana* to investigate the molecular functions of PaRXLR40. Our results demonstrate that PaRXLR40 suppresses immune responses and enhances pathogen colonization in *N. benthamiana*.

The ability of PaRXLR40 to suppress cell death triggered by different elicitors, including both apoplastic and cytoplasmic effectors, suggests that it may target core components of host defence signalling, rather than specific immune receptor pathways. This is consistent with findings from other *Phytophthora* RXLR effectors that modulate key host regulatory processes such as hormone signalling, vesicle trafficking, or nuclear functions to promote infection (Anderson *et al*., 2015; He *et al*., 2020).

To test the contribution of PaRXLR40 to virulence, we used RNAi to silence the effector during infection. Reduced *PaRXLR40* expression was associated with decreased *P. agathidicida* colonization in both *N. benthamiana* and *A. australis*, demonstrating its role in virulence across diverse plants. This supports the idea that PaRXLR40 targets conserved plant components to promote susceptibility. Given its crucial role in virulence, *PaRXLR40* represents a promising target for RNAi-based control strategies. Silencing pathogen effectors *in planta*, either through host-induced gene silencing or external application of dsRNA, could offer a novel approach to limit disease development (Sellamuthu *et al*., 2025).

Using a Y2H screen, we identified StARIA, an ARM/BTB-repeat protein, as a potential interactor of PaRXLR40. ARIA homologs have previously been described as positive regulators of ABA responses in *A. thaliana* (Kim *et al*., 2004). In *N. benthamiana*, we found that StARIA, like PaRXLR40, can suppress cell death triggered by both apoplastic and cytoplasmic cell death elicitors, suggesting a role in modulating core immune signalling pathways. Consistent with this, overexpression of StARIA in *N. benthamiana* enhanced *P. agathidicida* colonization, indicating that StARIA promotes susceptibility. Moreover, *StARIA* was transcriptionally induced during early stages of *P. agathidicida* infection in *N. benthamiana*, supporting the idea that the StARIA protein is exploited by the pathogen to facilitate host colonization and establishment. Together, these findings suggest that StARIA acts as a susceptibility factor. As such, it is unlikely that PaRXLR40 inhibits StARIA function but rather promotes it to suppress plant immunity and promote *P. agathidicida* virulence, as has been seen for other S factor targets (Boevink *et al*., 2016; Turnbull *et al*., 2017, 2019; He *et al*., 2020; Wang *et al*., 2023).

Supporting this, exogenous ABA application increased the susceptibility of both *N. benthamiana* and *A. australis* to infection by *P. agathidicida*. ABA is best known for its roles in regulating abiotic stress responses, particularly in drought and salinity tolerance (Sussmilch *et al*., 2017), but it also plays a complex and often antagonistic role in plant-pathogen interactions. In several plant-pathogen systems, elevated ABA levels have been associated with increased susceptibility to infection (Audenaert *et al*., 2002; Sánchez-Vallet *et al*., 2012; Ulferts *et al*., 2015; Sivakumaran *et al*., 2016), largely due to ABA’s capacity to antagonize key immune pathways, especially those mediated by SA, which is central to defence against biotrophic and hemibiotrophic pathogens (Singh & Roychoudhury, 2023). PaRXLR40 may promote ABA activity through its interaction with StARIA; however, whether PaRXLR40 directly manipulates ABA signalling or targets StARIA to indirectly influence downstream hormonal crosstalk remains to be determined.

Interestingly, we also observed that StARIA can form homodimers, a property mediated by its BTB domain and reliant on a conserved aspartate residue (Melnick *et al*., 2000). Mutation of a conserved aspartate residue (D542) within the BTB/POZ domain of StARIA disrupted dimerization and its ability to interact with PaRXLR40. This residue aligns with D146 in the BTB/POZ domain of protein POB1, in which mutation impairs dimerization, disrupts interaction with the E3 ligase PUB17, and abolishes suppression of cell death (Orosa *et al*., 2017). In contrast, the StARIA^D542A^ mutant retained its ability to suppress effector-triggered cell death and promote *P. agathidicida* infection in *N. benthamiana*. These findings suggest that, unlike POB1, StARIA may function independently of, or in addition to, BTB-mediated dimerization. This difference could reflect functional specialization within the BTB/POZ protein family or indicate that StARIA employs additional interaction surfaces or partners for its immune-suppressive activity.

To further characterize potential downstream targets of StARIA, we performed an additional Y2H screen and identified SOG1 as a specific interactor. *Arabidopsis* SOG1 is a NAC-domain transcription factor central to DNA damage response and oxidative stress signalling, and which positively regulates immunity (Yoshiyama *et al*., 2009; Yoshiyama *et al*., 2020). Overexpression of *N. benthamiana* SOG1 (NbSOG1) resulted in reduced lesion size upon *P. agathidicida* infection, suggesting enhanced resistance. NbSOG1 also interacted with StARIA but not PaRXLR40 in Y2H, pointing to a possible ARIA-SOG1 regulatory module involved in plant defence. The interaction between StARIA and NbSOG1, and their opposing effects on resistance, raises the possibility that PaRXLR40 may hijack the host ABA signalling pathway via interaction with StARIA to suppress immunity. Given that NbSOG1 also interacts with StARIA, it may compete for binding or modulate StARIA’s function, potentially acting as a resistance factor. However, further work is needed to confirm this interaction *in planta* and to determine whether PaRXLR40 or StARIA expression influences SOG1 protein levels during pathogen infection or upon protein delivery. Nevertheless, these findings point to a mechanism by which *P. agathidicida* exploits a regulatory hub between hormone signalling and stress responses to promote infection.

Overall, our findings support a model in which PaRXLR40 promotes *P. agathidicida* infection by suppressing host immunity and targeting an ARM/BTB-domain containing protein that may function in ABA-regulated defence. This study provides insight into how oomycete effectors exploit host signalling pathways, and highlights ARIA-like proteins as potential susceptibility factors in both host and model-plant systems. Future work should focus on defining the molecular mechanisms of PaRXLR40-StARIA interaction and the broader role of ABA in gymnosperm immunity.

## 5 Acknowledgements

The work was supported by the Tertiary Education Commission of New Zealand via Bioprotection Aotearoa grant number 39240 and by the Biotechnology and Biological Sciences Research Council – New Zealand partnering award BB/T020164/1.

Kauri germplasm was supplied by Scion with approval from Taoho Patuawa, representing the Te Roroa Iwi Trust. The project was undertaken with the cultural authority of Te Roroa, the mana whenua of the Waipoua region.

## 7 Author contributions

MT, YG, HM, ELB, REB, PCB, PRJB and CHM planned and designed the research. MT, YG, HM and ELB performed the experiments. MT, YG, REB and CHM wrote the manuscript. All authors reviewed the manuscript and approved it for publication.

## 8 Data availability

All data can be found in this manuscript and in its Supplementary Information.

## 10 Supporting Information

**Fig. S1.**
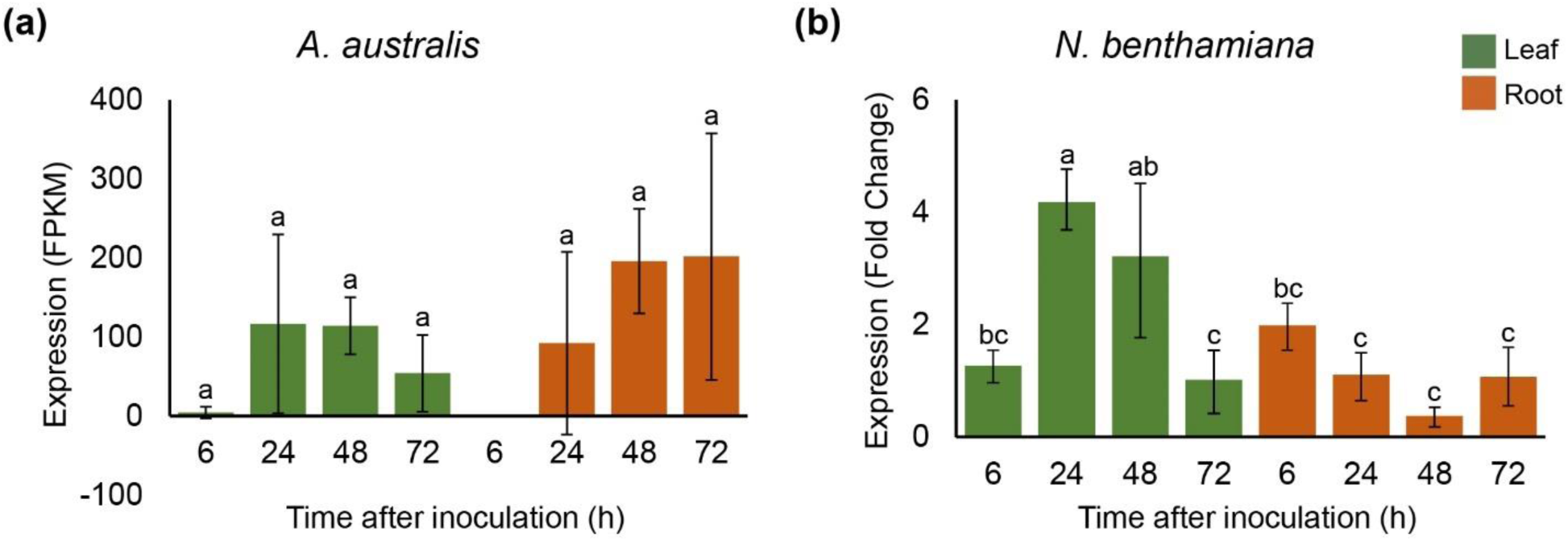
*PaRXLR40* expression during infection of *Agathis australis* and *Nicotiana benthamiana*. (a) Fragments Per Kilobase per Million (FPKM) values of *PaRXLR40* were obtained from transcriptomic data of *P. agathidicida* infecting *A. australis* at 6, 24, 48, and 72 h post inoculation (hpi) in leaves (green) and roots (brown) (Cox et al., 2022). (b) Expression (Fold Change) of *PaRXLR40* in response to *P. agathidicida* inoculation of *N. benthamiana* leaves (green) and roots (orange) at 6, 24, 48 and 72 hpi. Transcript levels were normalized to the reference genes *PaActin* and *Paβ-tubulin*. Means and standard errors were calculated from three biological replicates. Bars represent mean ± SD; different letters indicate statistically significant differences (*P*<0.05, Tukey’s test).

**Fig. S2.**
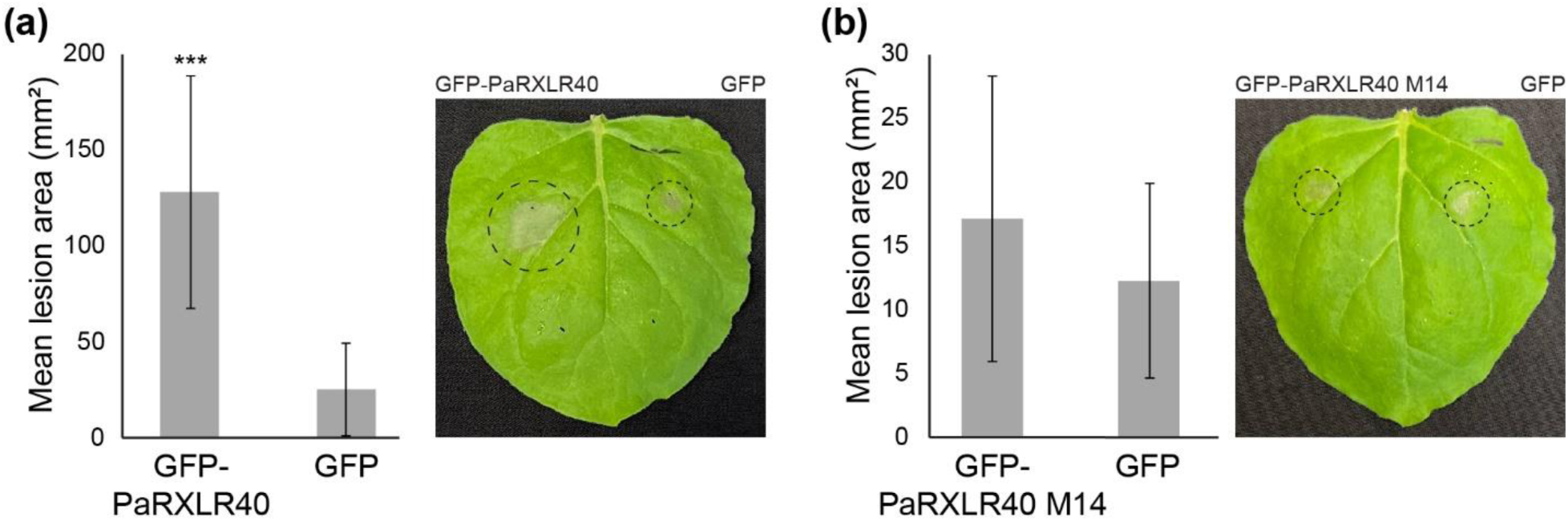
PaRXLR40 enhances *Phytophthora infestans* infection in *Nicotiana benthamiana*. Overexpression of (a) GFP-PaRXLR40 in *N. benthamiana* enhances *P. infestans* leaf colonization, while overexpression of (b) PaRXLR40 mutant 14 (M14) shows similar levels of infection to the GFP control. *Agrobacterium tumefaciens* carrying expression vectors for each protein were infiltrated in opposing leaf segments of *N*. *benthamiana*. Leaves were then inoculated with *P*. *infestans* at 1 day after infiltration with *A. tumefaciens*. Photos and measurements of lesion area (mm^2^) were taken 4 days post inoculation. Means and standard errors were calculated from four biological replicates. ***, *P*<0.001 using Student’s *t*-test.

**Fig. S3.**
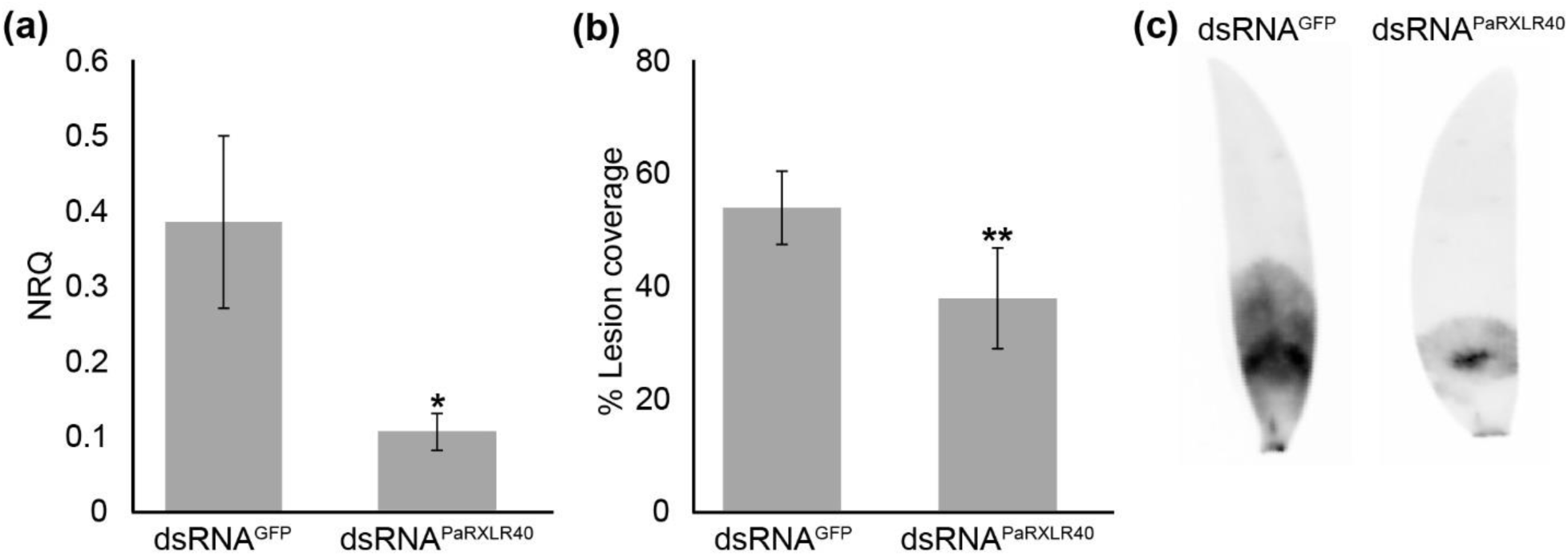
Silencing of *PaRXLR40* reduces *Phytophthora agathidicida* lesion size in *Agathis australis*. (a) Relative expression of *PaRXLR40* in *P. agathidicida*-infected leaves of *A. australis* at 48 hours post inoculation, following prior treatment with dsRNA^PaRXLR40^ or dsRNA^GFP^. Here, dsRNA leaf treatments were carried out 24 h before pathogen inoculation. Normalized relative quantity (NRQ) values represent normalized relative quantification of *PaRXLR40* transcript levels, calculated using *PaActin* and *Paβ-tubulin* as reference genes. Means and standard errors were calculated from three biological replicates. (b) Quantification of disease symptoms in *A. australis* as percentage (%) lesion coverage. Lesion coverage was quantified as the percentage of the leaf area affected by disease, calculated as the ratio between lesion length and total leaf length for each sample. Means and standard errors were calculated from eight biological replicates. **, *P*<0.01 using Student’s *t*-test. (c) Representative infrared images of disease symptoms at 5 days post inoculation.

**Fig. S4.**
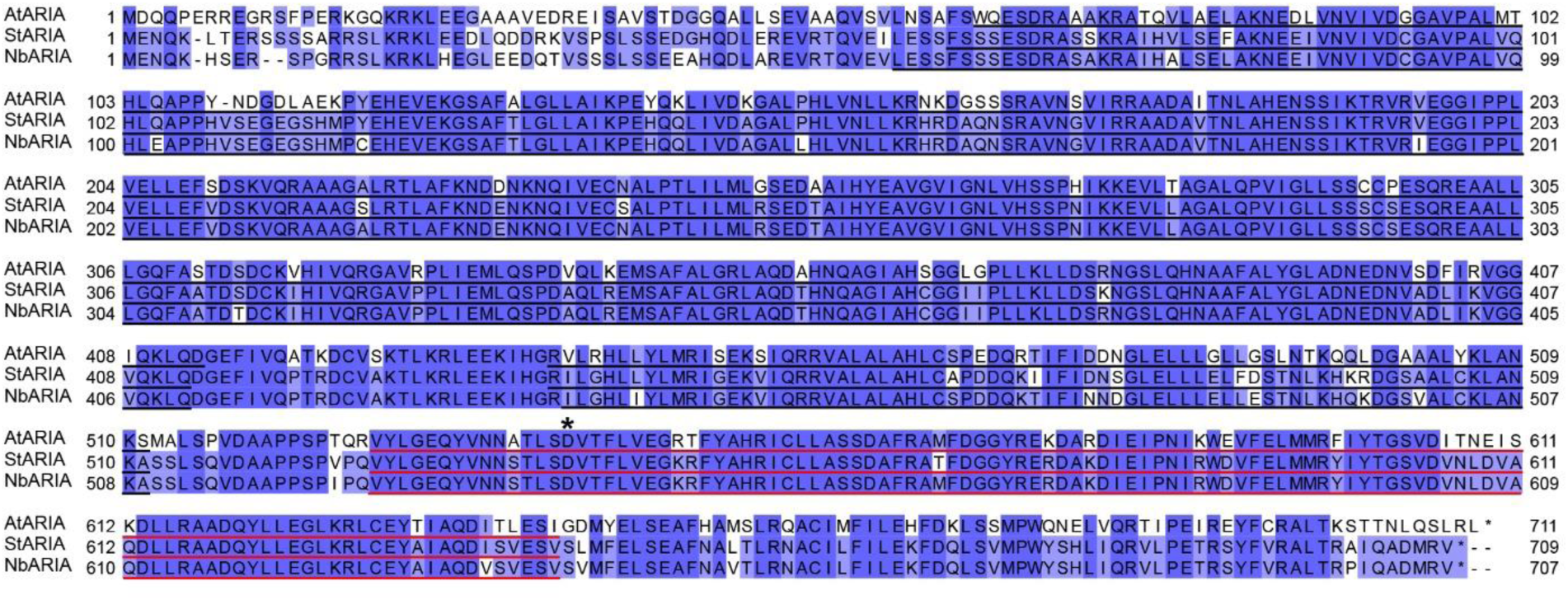
Sequence conservation of ARIA orthologues. Multiple sequence alignment of ARIA orthologs from *Arabidopsis thaliana* (AtARIA - NP_850852.1), *Nicotiana benthamiana* (NbARIA - Niben101Scf02021g02009.1), and *Solanum tuberosum* (StARIA - PGSC0003DMP400022811). The alignment was generated using Clustal Ω (Sievers et al., 2011), with blue shading indicating the degree of amino acid sequence conservation. ARM repeat regions are underlined in black, and the BTB/POZ domain is highlighted in red, according to InterProScan predictions (Blum et al., 2025). The conserved aspartate residue (D542 in StARIA) mutated in this study is indicated by an asterisk (*).

**Fig. S5.**
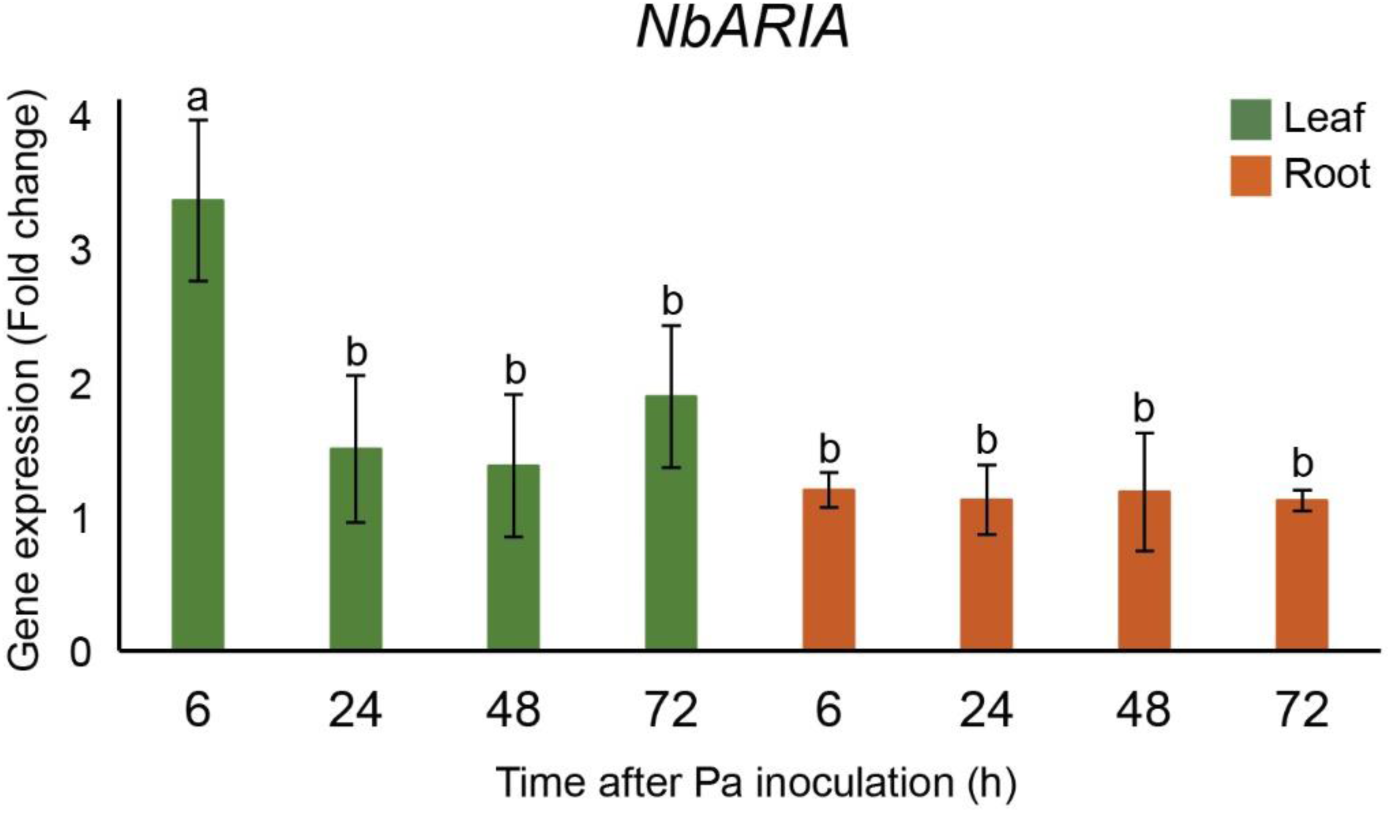
*NbARIA* is upregulated in *Nicotiana benthamiana* leaves during early stages of *Phytophthora agathidicida* infection. Expression (fold change) of *NbARIA* in response to *P. agathidicida* inoculation of *N. benthamiana* leaves (green) and roots (orange) at different time points (6, 24, 48 and 72 h). Transcript levels were normalized to the reference gene *NbActin*. Means and standard errors were calculated from three biological replicates. Means with different letters are significantly different from each other as determined by the Tukey test, with 95% confidence level.

**Fig. S6.**
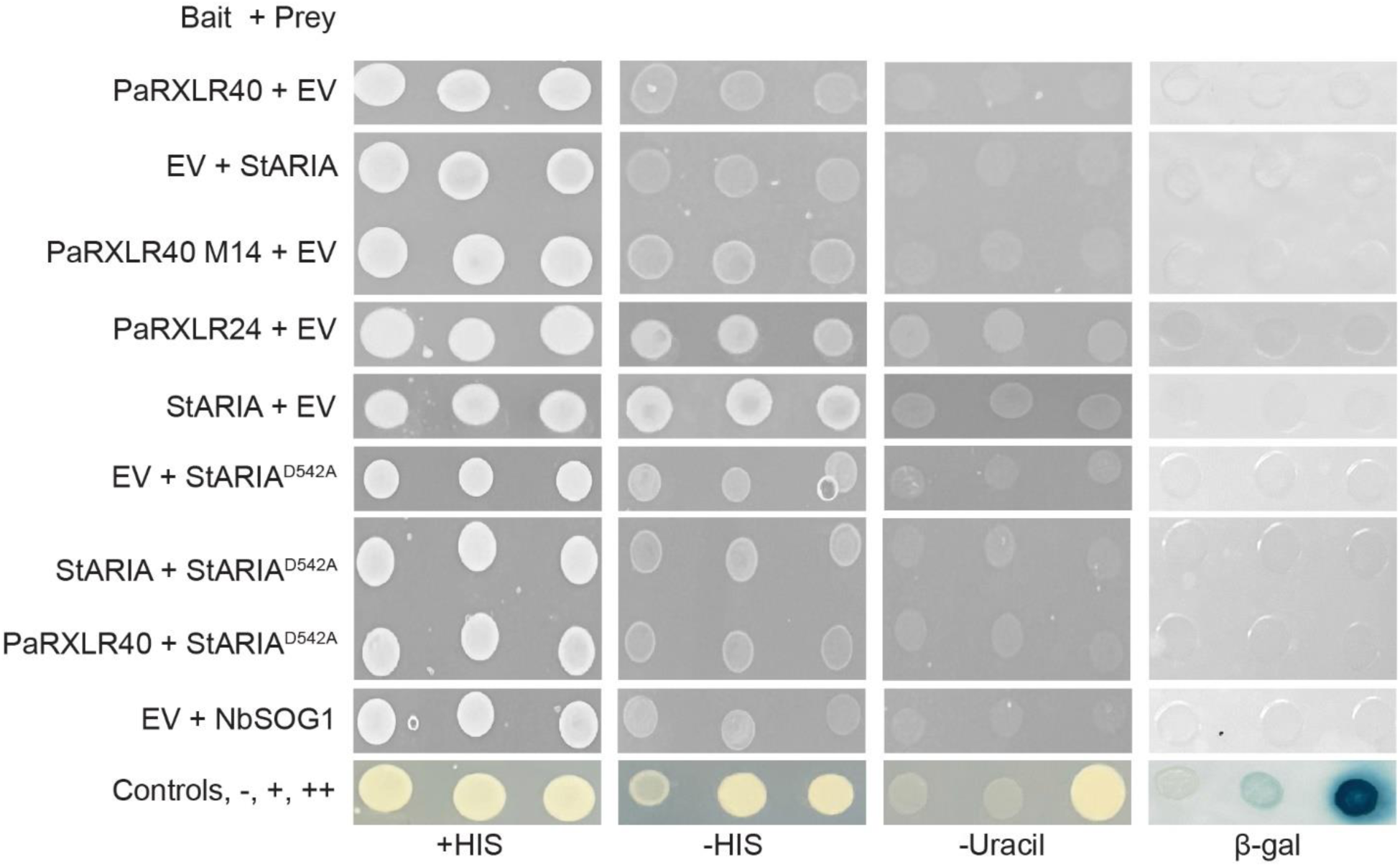
Yeast two-hybrid analysis of the PaRXLR40 and StARIA interaction. Yeast strain MaV203 was co-transformed with the indicated bait (PaRXLR40, PaRXLR40 mutant, PaRXLR24, or empty vector (EV)) and prey (StARIA, StARIA or empty vector) constructs and plated on media containing histidine (+HIS; growth control) or lacking histidine (-HIS) or uracil (-Uracil), as well as tested for β-galactosidase (β-gal) activity. None of the individual constructs exhibited reporter activity when co-expressed with the EV, confirming the absence of autoactivation. Truncated constructs of StARIA (BTB and ARM domains) were also tested with PaRXLR40 but did not show interaction. Yeast controls were: −, no interaction; +, weak interaction; ++, strong interaction.

**Fig. S7.**
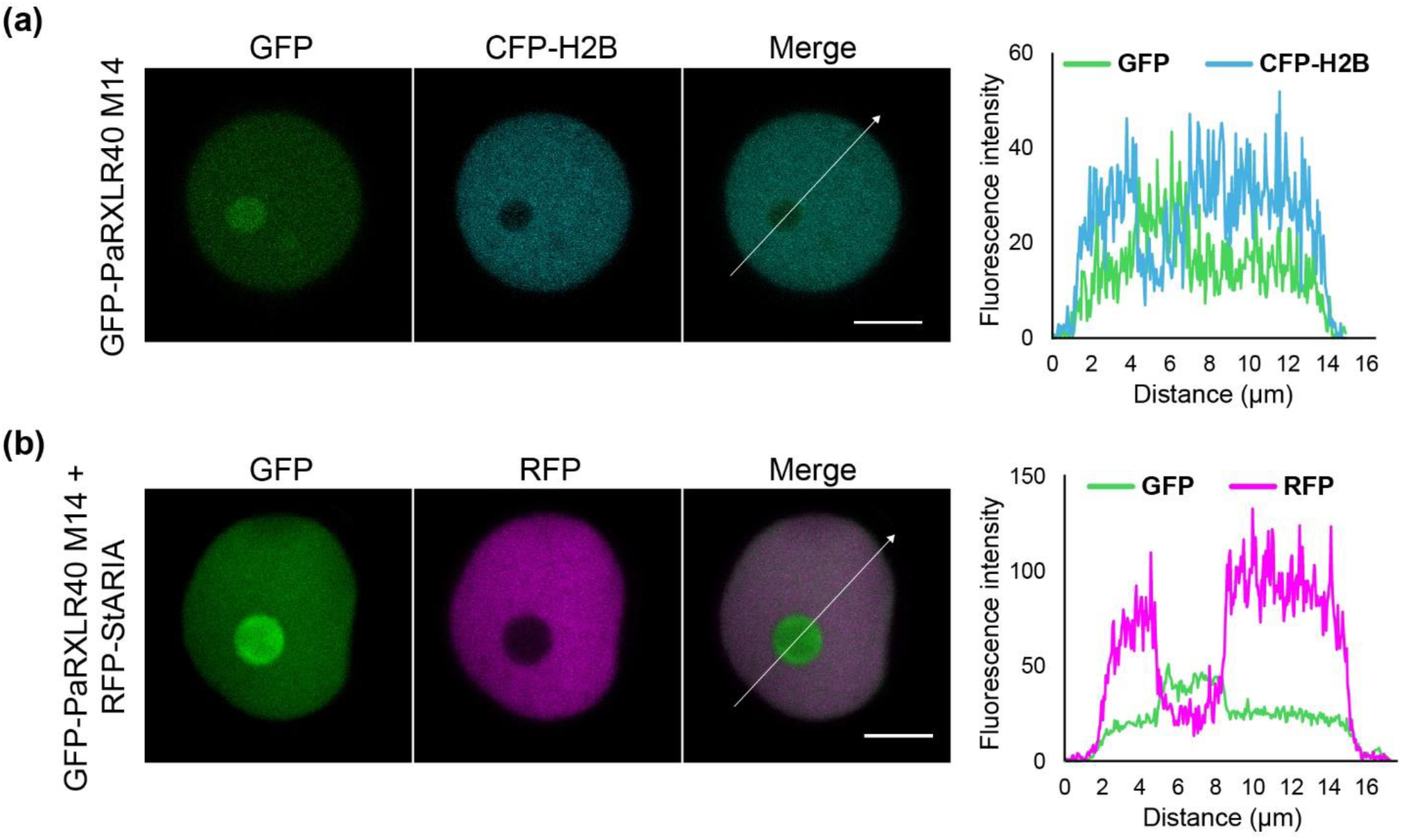
PaRXLR40 mutant and StARIA co-localize in the nucleus in *Nicotiana benthamiana*. (a) Representative confocal images of *N. benthamiana* epidermal cells transiently expressing GFP-PaRXLR40 M14 together with CFP-H2B (nuclear marker). GFP-PaRXLR40 M14 localizes to the nucleus and nucleolus. (b) Co-expression of GFP-PaRXLR40 M14 and RFP-StARIA shows overlapping nuclear localization without altered distribution patterns. White arrows indicate the transects used to generate the fluorescence intensity plots shown to the right of each image set. The X-axis of each plot represents the distance (in µm) along the corresponding white arrow. Scale bars = 5 µm.

**Fig. S8.**
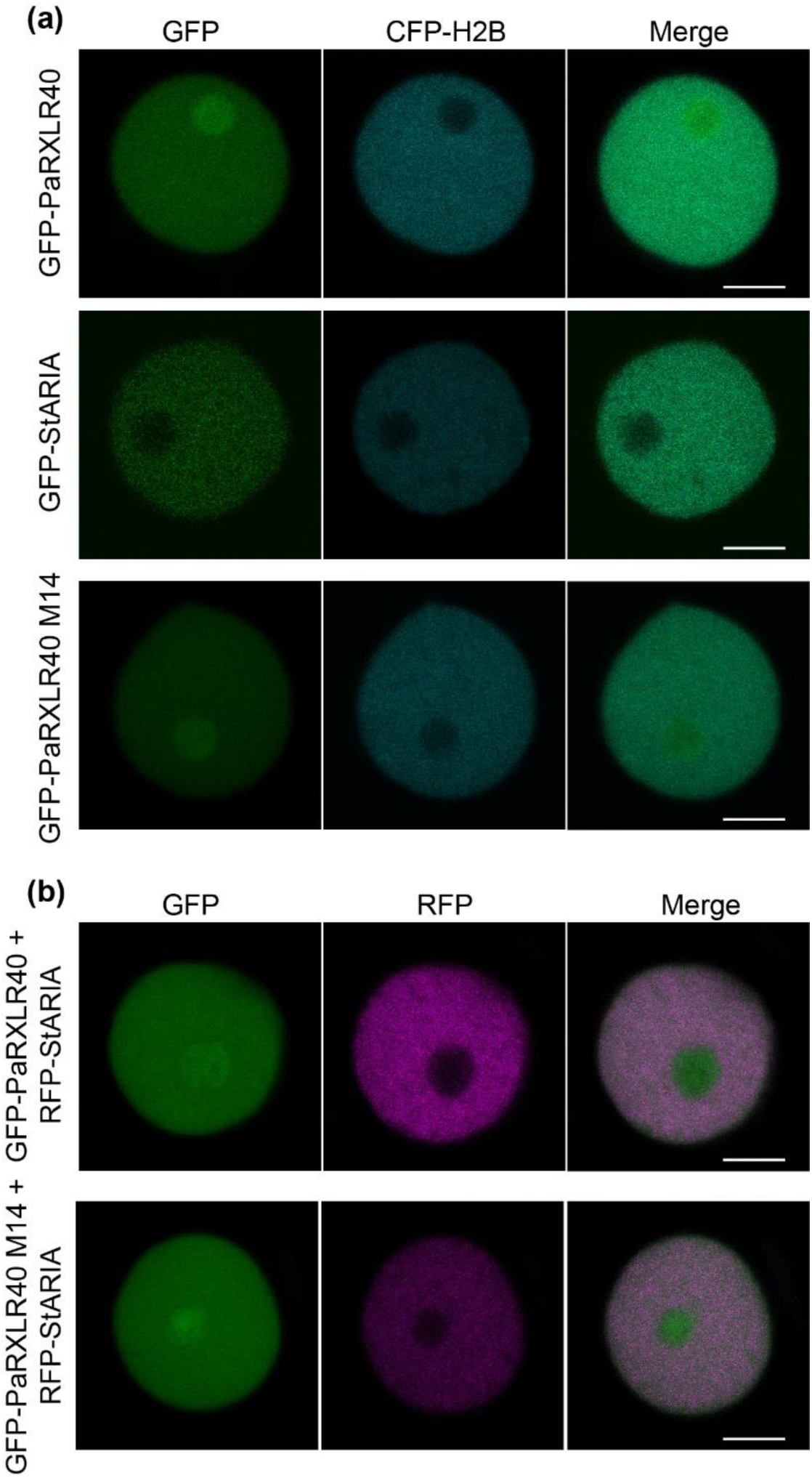
PaRXLR40, PaRXLR40 mutant and StARIA co-localize to the nucleus of *Nicotiana benthamiana*. (a) Representative confocal images of *N. benthamiana* epidermal cells transiently expressing GFP-PaRXLR40, GFP-PaRXLR40 M14 or GFP-StARIA, together with CFP-H2B (nuclear marker; CFP fused to histone H2B). GFP-PaRXLR40 and GFP-PaRXLR40 M14 localize to the nucleus and show enrichment in the nucleolus, while GFP-StARIA localizes to the nucleoplasm but is excluded from the nucleolus. (b) Co-expression of GFP-PaRXLR40 and RFP-StARIA, and GFP-PaRXLR40 M14 and RFP-StARIA show overlapping nuclear localization without altered distribution patterns. Scale bars = 5 µm.

**Fig. S9.**
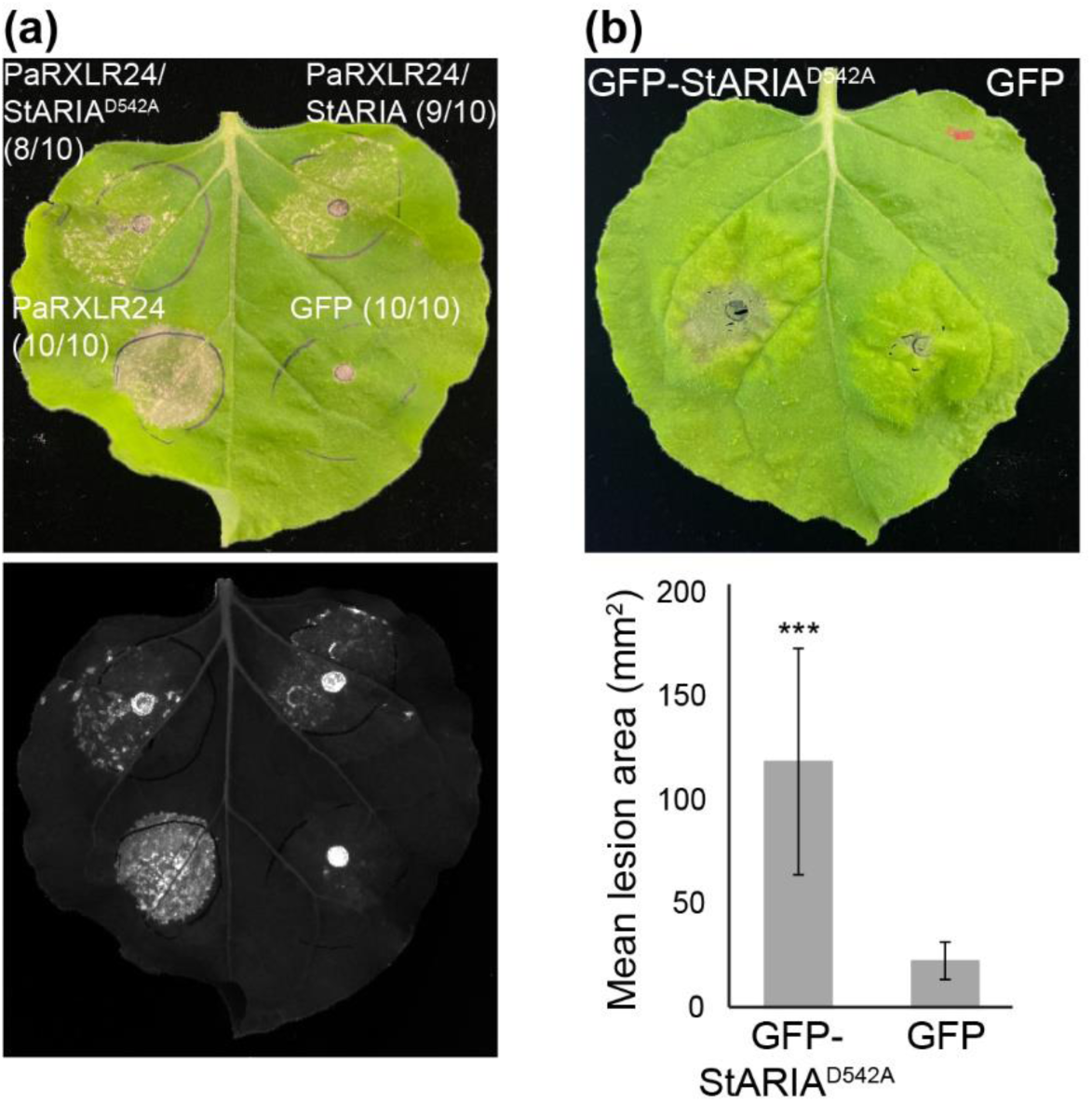
StARIA^D542A^ suppresses cell death and enhances *Phytophthora agathidicida* infection in *Nicotiana benthamiana*. (a) Suppression of PaRXLR24-triggered cell death by the StARIA^D542A^ mutant in *N. benthamiana*. *Agrobacterium tumefaciens* carrying an expression vector for PaRXLR24 was infiltrated 1 day after infiltration (dai) of *A. tumefaciens* carrying expression vectors for StARIA or StARIA^D542A^. Photographs with visible (top) and UV (bottom) light were taken 6 dai of *A. tumefaciens* carrying the PaRXLR24 expression vector. Representative images are shown from three independent experiments. Numbers in parentheses indicate the number of times the response was observed (left) out of the number of times the agroinfiltration was performed (right). (b) Overexpression of GFP-S StARIA^D542A^ in *N. benthamiana* enhances *P. agathidicida* leaf colonization. *A. tumefaciens* carrying each protein were infiltrated opposing leaf segments of *N. benthamiana*. Leaves were then inoculated with *P. agathidicida* at 1 dai with *A. tumefaciens*. Photos and measurements of lesion area (mm^2^) were taken at 4 days post inoculation with *A. tumefaciens*. Means and standard errors were calculated from four biological replicates. ***, P<0.001 using Student’s t-test.

**Fig. S10.**
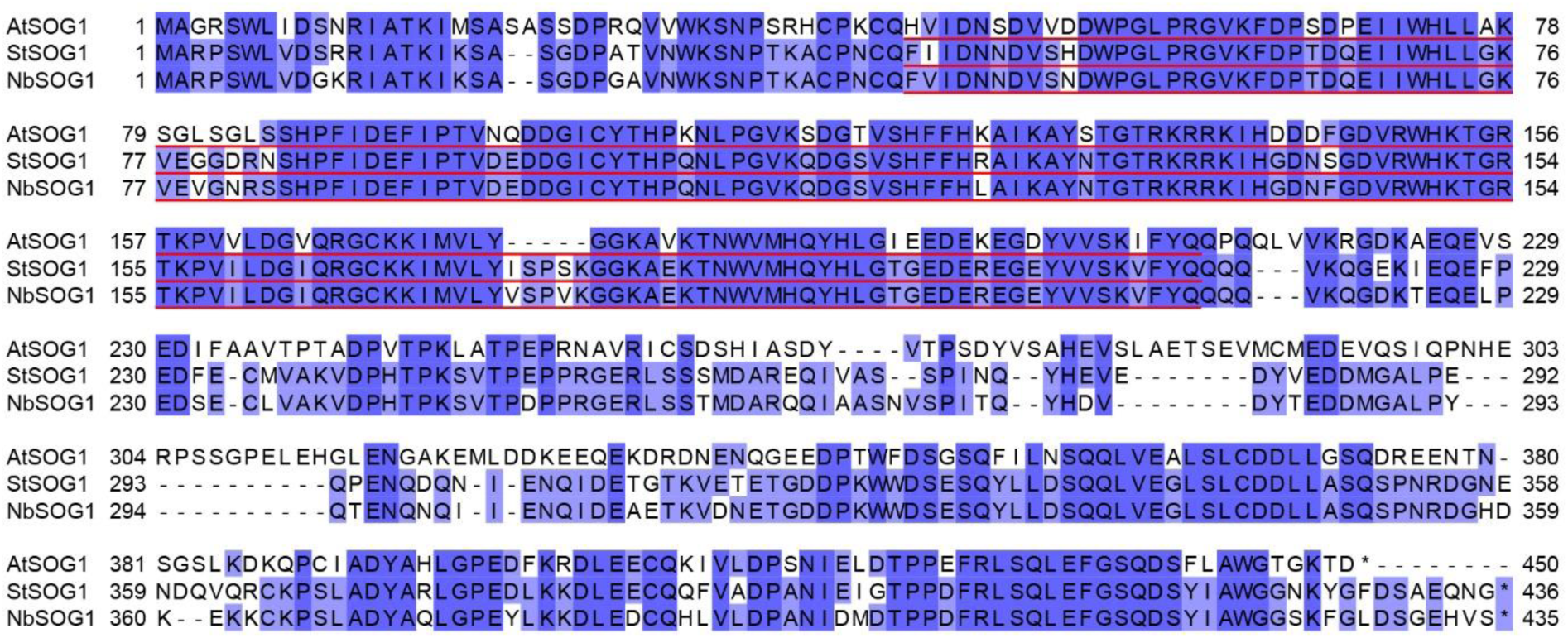
Sequence conservation of SOG1 orthologues. Multiple sequence alignment of SOG1 orthologs from *Arabidopsis thaliana* (AtSOG1 - NP_564238.2), *Nicotiana benthamiana* (NbSOG1 - Niben101Scf01627g02012.1), and *Solanum tuberosum* (StSOG1 - PGSC0003DMP400001544). The alignment was generated using Clustal Ω (Sievers et al., 2011), with blue shading indicating the degree of amino acid sequence conservation. The NAC domain is highlighted in red, according to InterProScan predictions (Blum et al., 2025).

